# Investigating the axoneme CCDC40 protein reveals new insights in trypanosome morphogenesis and division

**DOI:** 10.64898/2026.05.22.727155

**Authors:** Christine Girard-Blanc, Thierry Blisnick, Vincent Louvel, Paul Guichard, Virginie Hamel, Philippe Bastin

## Abstract

Cilia and flagella contribute to cell morphogenesis in multiple organisms. In the parasite *Trypanosoma brucei*, the flagellum is attached along the length of the cell body and acts as a guide for cell division, while its motility function is required for the completion of cytokinesis. To tease apart the contributions of flagellum length and motility to trypanosome morphogenesis, we investigated the coiled-coil containing domain 40 (CCDC40 or FAP172) protein. Iterative Ultrastructure Expansion Microscopy (iU-ExM) revealed that CCDC40 is associated to the 96-nm repeats of the trypanosome axoneme. CCDC40 depletion by RNAi leads to loss of components from the dynein regulatory complex, inner dynein arms and radial spokes, resulting in disconnected microtubule doublets and disorganised axoneme structure, abrogating motility and resulting in flagella and cell bodies 2-3 times shorter than normal. We show for the first time that this short flagellum phenotype is associated to slower tubulin incorporation and premature acquisition of the maturation marker FLAM8 but not of the locking protein CEP164C. Surprisingly, these short and immotile trypanosomes grow and divide normally. We discuss the significance of these observations for trypanosome morphogenesis and division.

## Introduction

In addition to their well-known motility and sensory functions, cilia and flagella are important drivers of cell morphogenesis in multiple organisms. In animal cells, photoreceptors and spermatozoa are typical examples where the cellular architecture is built around these organelles (Salinas et al., 2017; San Agustin et al., 2015). This is also true in flagellated protists, most of them maintaining their existing flagella during cell division (Gillott and Triemer, 1978; Kato et al., 2000; Konupkova et al., 2025; Nohynkova et al., 2006; Sherwin and Gull, 1989), in contrast to animal cells. The flagellum is a key actor of cell morphogenesis in the parasite *Trypanosoma brucei* (Kohl et al., 2003), which is responsible for sleeping sickness in Central Africa. This organism is an excellent model for cell biology and evolution (Lukes et al., 2023), in particular for the study of cilia and flagella (Vincensini et al., 2011). It is transmitted by the bite of the tsetse fly and encounters multiple environments during its life cycle, accompanied by specific biochemical and morphological modifications, including in flagellum length that can vary from 3 to 30 µm (Rotureau et al., 2011) (Matthews, 2005).

Trypanosomes belong to the Kinetoplastea class characterised by the presence of a single mitochondrion that contains the kinetoplast (mitochondrial DNA), whose replication is coupled to that of the basal body of the flagellum (Robinson and Gull, 1991) and whose position varies between life cycle stages (Van Den Abbeele et al., 1999). In the procyclic and bloodstream stages, trypanosomes exhibit an elongated shape, with a cell body length ranging between 20 and 25 µm for a width of 3-5 µm. A corset of tightly-spaced subpellicular microtubules defines cell shape (Sherwin and Gull, 1989). The flagellum emerges via a flagellar pocket found at the posterior end and is attached to the cell body for most of its length, with the exception of its last few µm (Sherwin and Gull, 1989). This is called the trypomastigote configuration while other stages adopt the epimastigote, characterised by the positioning of base of the flagellum between the nucleus and the anterior end of the cell (Hoare, 1966). Attachment is mediated by a structure called the flagellum attachment zone or FAZ, which is constituted of a filament present on the cell body side found in a gap between two corset microtubules and is itself linked to the flagellum via a complex series of proteins (review in (Sunter and Gull, 2016)).

During the cell cycle, neither the microtubule corset nor the flagellum and its associated structures depolymerise, posing constraints on cell replication, which can be considered as of three cycles: the nucleus, the kinetoplast (mitochondrial genome) and the cytoskeletal elements including the flagellum (Wheeler et al., 2019). A new flagellum emerges from the posterior end, associated to a new FAZ, and the tip of the flagellum follows the existing one during elongation guided by a flagella connector in the procyclic stage (Briggs et al., 2004; Moreira-Leite et al., 2001) and buried withing a specific groove in the bloodstream stage (Hughes et al., 2013). The nucleus goes through S-phase and mitosis before cytokinesis is initiated from the anterior end of the cell, having to navigate through the microtubule corset along the long axis of the cell towards the posterior end (Fig. S1).

Flagella are major contributors to trypanosome morphogenesis and division. The length of the new flagellum can be reduced by inducible RNAi knockdown of proteins belonging to the intraflagellar transport (IFT) complex, the machinery responsible for flagellum construction. As a result, shorter cells are produced, with a direct correlation between flagellum and cell length (Kohl et al., 2003). This is in agreement with the hypothesis that the tip of the new flagellum controls the initiation point of cytokinesis, probably by positioning the anterior end of the FAZ (Robinson et al., 1995; Zhou et al., 2011). At later time points of RNAi induction targeting IFT components, short non-flagellated cells are produced and fail to divide (Kohl et al., 2003). Flagellum attachement is also critical as shown upon depletion of FAZ components which leads to cytokinesis failure (Sun et al., 2013; Vaughan et al., 2008). Flagella remain motile throughout the cell cycle and in the final stage of cell division, they point in opposite directions (Wheeler et al., 2013). Inhibition of flagellum beating blocks cell division (Branche et al., 2006; Broadhead et al., 2006; Ralston et al., 2006), suggesting a mechanical contribution of flagellum motility to the completion of cell cleavage.

To further define the role of the flagellum in cell morphogenesis, we aimed to produce trypanosomes with short and immotile flagella. We took advantage of the molecular and structural conservation of the trypanosome axoneme to browse the literature in search of candidate mutations that could result in such flagella. Our attention was caught by CCDC40 (coiled-coil containing protein 40), also termed FAP172. Biochemical and structural data revealed that CCDC40 dimerises with CCDC39, another coiled-coil containing protein, to form an elongated filament along the length of the A-microtubule of the axoneme doublet and is the ruler responsible for the 96-nm repeats (Oda et al., 2014). Absence of CCDC40 (or of its partner CCDC39) in human patients, in mouse models or in the green alga *Chlamydomonas* has severe consequences on flagellum anatomy as it leads to loss of the dynein-regulatory complex, of inner dynein arms and of radial spokes (Antony et al., 2013; Aprea et al., 2023; Becker-Heck et al., 2011; Blanchon et al., 2012; Brody et al., 2025; Merveille et al., 2011). As a result, microtubule doublets appear disconnected, and cilia fail to beat but also are shorter than usual, a phenotype that so far remains unexplained. CCDC40 is conserved in trypanosomes and has been identified in several proteomic studies of purified trypanosome flagella (Broadhead et al., 2006; Subota et al., 2014). Recent cryo-electron microscopy data of *T. brucei* flagellar microtubules demonstrated a similar organisation of the 96-nm repeats as in other eukaryotes, with the conserved CCDC39/CCDC40 dimer (Xia et al., 2025).

We have therefore tagged the trypanosome CCDC40 protein and are showing its precise distribution by Iterative Ultrastructure Expansion Microscopy (iU-ExM). Knocking-down CCDC40 by RNAi has severe impact on the structure and the motility of the flagellum, but also its length, which is 2-3 times shorter than in control cells. We provide for the first time a potential explanation for this phenotype by revealing that tubulin incorporation is reduced by 2-fold while the FLAM8 maturation marker, but not the locking protein CEP164C, is acquired earlier. In agreement with previous data, assembly of a shorter flagellum results in shorter cells. Despite the absence of flagellum beating, these short trypanosomes grow and divide normally. They possess a shorter FAZ that lines up the short flagellum, which adheres properly to the cell body.

## Results

### CCDC40 localises to the 96-nm repeats of the trypanosome axoneme

The *T. brucei CCDC40* gene (Tb927.11.11250) encodes a protein of 872 aa with a predicted molecular weight of 101 kDa that contains numerous coiled-coil domains. It exhibits 25-35% identity with the corresponding protein in human and *Chlamydomonas*. CCDC40 was tagged at its C-terminal domain with a GFP and 6 copies of the Ty-1 tag using the pPOTv6 system which allows endogenous tagging (Dean et al., 2015). Trypanosomes are diploid, so the second allele was tagged, this time with a mNeonGreen (mNG) and 6 copies of the Ty-1 tag, by using consecutive nucleofection with two different drug resistance markers (Fig. S2A). The double tagged cell line therefore expresses CCDC40::GFP and CCDC40::mNG. Since both copies contain Ty-1 tags, the fusion proteins will be refered to as CCDC40^Ty^ for simplicity when discussing immunofluorescence experiments performed with the double-tagged cell line. PCR analysis confirmed proper tagging of one or two alleles (Fig. S2A-B) while western blot analysis with the BB2 antibody that detects the Ty-1 tag showed the fusion proteins migrated according to their expected molecular weight around 130 kDa (Fig. S2C). Since GFP and mNG have a similar molecular weight, the two fusion proteins cannot be discriminated and so tagging both alleles produced a stronger BB2 signal (Fig. S2C). Live imaging confirmed expression of the CCDC40::GFP fusion protein and association to the flagellar compartment (Fig. 1A, example of single-allele tagging). It did not visibly alter neither cell growth nor motility and morphology, suggesting that the fusion protein is functional. Cell fractionation revealed that most of the protein was present in the cytoskeletal pellet, with only a low amount found as soluble material (Fig. 1B), in agreement with the expected tight association of CCDC40 to the axoneme.

**Figure 1.**
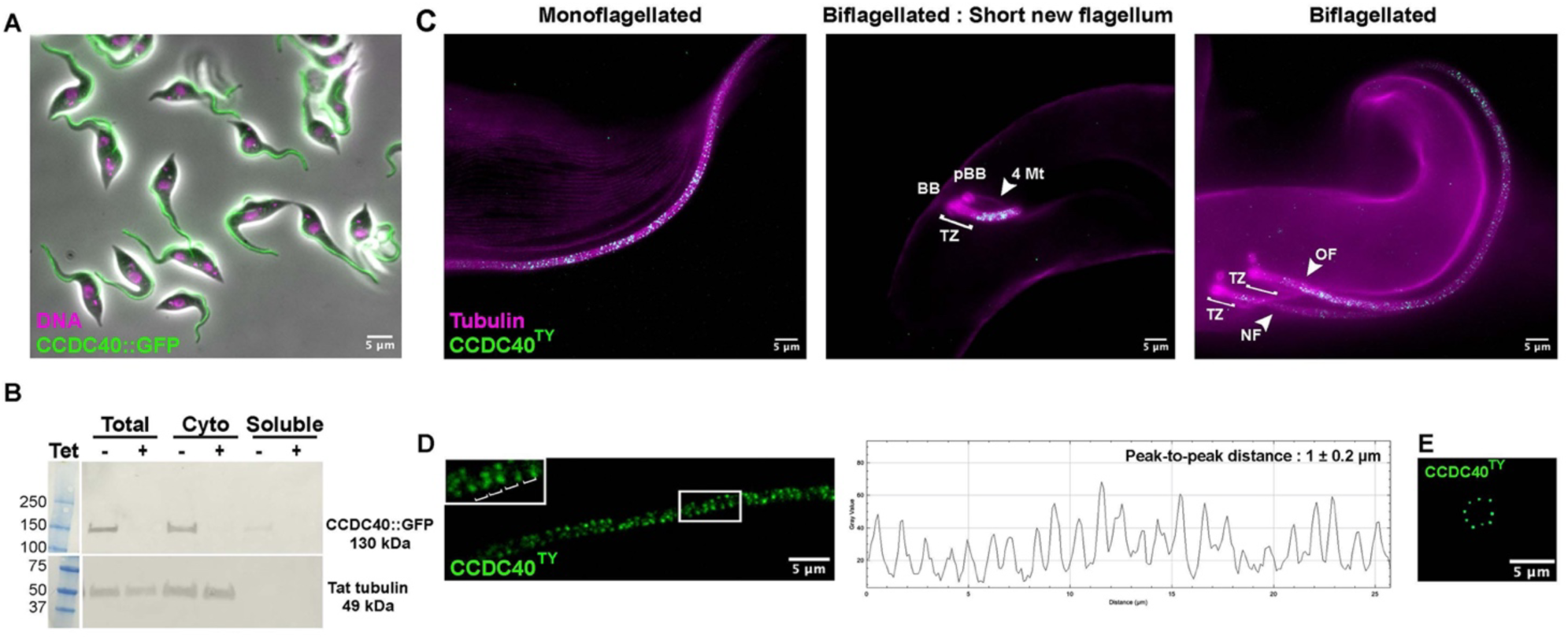
CCDC40 localises to the axonemal 96-nm repeats. (A) Live *CCDC40^RNAi1-500^* cells expressing CCDC40::GFP (single-allele tagged, green) grown in the absence of tetracycline were washed, stained with Hoechst (magenta) and observed directly. The fusion protein is present all along the length of both old and new flagella. (B) Western blot of samples from the *CCDC40^RNAi1-500^* cell line expressing CCDC40::GFP in absence (-) or presence (+) of tetracycline. The membrane was cut in two and the top part was probed with BB2, recognising the Ty1 epitope tags present on the CCDC40::GFP fusion protein while the bottom panel was revealed with TAT-1, which detects alpha-tubulin and was used as a loading and fractionation control. Molecular weight markers are indicated on the left. Most CCDC40::GFP material remains associated with the cytoskeletal fraction, with only minimal amounts detected in the detergent-soluble fraction. All CCDC40::GFP bands disappear after addition of tetracycline for 48 hours, demonstrating the efficiency of RNAi. (C-D) Iterative expansion microscopy (iU-ExM) of cells expressing CCDC40::GFP and CCDC40::mNG (so both alleles tagged) here called CCDC40^Ty^ stained with BB2 (green) and minibodies against alpha- and beta-tubulin (magenta), which label the subpellicular microtubules and the flagella, including the basal body (BB) and the pro-basal body (pBB). The quartet microtubules (4Mt) are also visible. (C) On a side-view, the CCDC40^Ty^ signal appears as dots associated to flagellar microtubules and regularly distributed along the length of the axoneme. Representative examples of a uniflagellated cell and two biflagellated cells, with the new flagellum at early or later stages of elongation. CCDC40^Ty^ is missing from the basal bodies (BB and pro-basal body, pBB) and the transition zone (TZ), in agreement with the absence of dynein arms in these areas. For the cell in the middle panel, the focus is made on the short new flagellum while the mature flagellum is out of the plane, hence not visible. (D) Lateral view of the CCDC40::GFP signal (green) showing its repetitive pattern. The graph shows signal intensity after one-dimensional projection of this flagellum. Peaks are spaced by 1.0 ± 0.2 µm corresponding to an actual separation of ∼91 nm taking into account the 11-fold expansion. (E) A section through the flagellum shows the presence of nine CCDC40^Ty^ spots corresponding to the expected presence on all nine doublet microtubules (green).

To define the exact position of the CCDC40 fusion proteins within the flagellum, we used iterative Ultrastructure Expansion Microscopy (iU-ExM), which allows up to 16-fold expansion of the sample, reaching the resolution necessary to visualise the complex axonemal architecture (Louvel et al., 2023). Cells were co-stained with the BB2 anti-Ty1 antibody (Bastin et al., 1996) and with minibodies recognising alpha- and beta-tubulin (Araujo Alves et al., 2025). Because of the axoneme cylindrical shape, tubulin staining revealed a maximum of 4 out of its 9 doublets when viewed from the side (magenta signal, Fig. 1C). The CCDC40^Ty^ signal appeared as dots repeating at regular intervals along each microtubule doublet (green signal, Fig. 1C). As expected, the signal looked less bright in cells where a single allele was tagged (Fig. S2D), due to competition with the untagged protein, compared when both alleles were modified (Fig. S2E). The signal was found in both mature and growing flagella (Fig. 1C), as soon as the new flagellum extended beyond the transition zone (Fig. 1C, central panel for a very short flagellum and right panel for a longer one). CCDC40^Ty^ is absent from the basal bodies and from the transition zone (Fig. 1C, central and right panels), in agreement with the absence of dynein arms in this area. Virtually no signal could be detected on subpellicular microtubules either. Higher magnification highlighted the repetitive nature of the CCDC40^Ty^ signal along the length of the axoneme (Fig. 1D). Plotting signal intensity after one-dimensional projection showed a repeat pattern of 1.0 ± 0.27 µm (n=90 from 5 different flagella), which when taking into account the expansion factor of 11x in that particular experiment, corresponds to an actual spacing of ∼91 nm (Fig. 1E), quite close to expectations from cryo-electron microscopy data (Xia et al., 2025). Finally, by sectioning the gel, transversal views of the axoneme were visible and the CCDC40^Ty^ fusion protein displayed nine spots as expected (Fig. 1F). These data demonstrate that the CCDC40^Ty^ fusion protein localises to its expected location as proposed by cryo-electron microscopy analysis (Xia et al., 2025) and is fully functional since it can replace the two copies of the wild-type allele product without visibly impacting trypanosome behaviour, at least in culture.

### Silencing of CCDC40 disrupts axonemal organisation

The expression of CCDC40 was knocked down by tetracycline-inducible expression of double-stranded RNA. Two different 500 bp segments were selected covering either residues 1 to 500 or residues 1100 to 1599 and cloned in a vector ensuring tetracycline-inducible expression of double-stranded RNA. These plasmids were introduced in the 29-13 recipient cell line (Wirtz et al., 1999), resulting in cell lines *CCDC40^RNAi1-500^* and *CCDC40^RNAi1100-1599^*. Since both knockdowns yielded similar phenotypes, the term *CCDC40^RNAi^*will be used throughout the manuscript for the sake of readibility. Details of which cell line was used is given in figure legends. To allow direct monitoring of the silencing efficiency, RNAi was performed in cell lines expressing endogenously tagged CCDC40::GFP as reporter. Western blot investigation showed a rapid and potent reduction in the amount of CCDC40::GFP after addition of tetracycline and production of *CCDC40* dsRNA in both single and double allele tagged cell lines (Fig. 1B and Fig. S2C). Similarly, live imaging revealed a strong reduction of the fluorescent signal after 48 hours in RNAi conditions (Fig. 2A-B). To monitor CCDC40 depletion, we fixed cells and performed immunofluorescence assay using BB2 to detect the CCDC40::GFP fusion protein and the axonemal marker mAb25 that recognises the SAXO protein (Dacheux et al., 2012), a microtubule inner protein (Xia et al., 2025). In control conditions, CCDC40::GFP was found equally in both old and growing flagella (Fig. 2C). Since RNAi targets only mRNAs, proteins assembled before knockdown onset were still present at the early phase (18h in tetracycline) and were detected associated to the old but not the new flagellum (Fig. 2D). At more advanced time points, both flagella became deprived of CCDC40::GFP (Fig 2E). CCDC40 is predicted to associate with CCDC39 and to constitute the anchor for the inner dynein arms (IDA), the dynein regulatory complex (DRC) and the radial spokes (RS)(Brody et al., 2025; Walton et al., 2023; Xia et al., 2025). To evaluate the impact of CCDC40 depletion on these structures, a component of each of them was endogenously tagged in the *CCDC40^RNAi^* cell line as described above (Dean et al., 2015). Western blotting analysis revealed a marked reduction in the flagellar amounts of CCDC39 (CCDC40-binding partner), IDA1 (inner dynein arm component), DRC2 (member of the dynein regulatory complex) and RSP3 (protein of the radial spoke) in the absence of CCDC40 (Fig. 2F).

**Figure 2.**
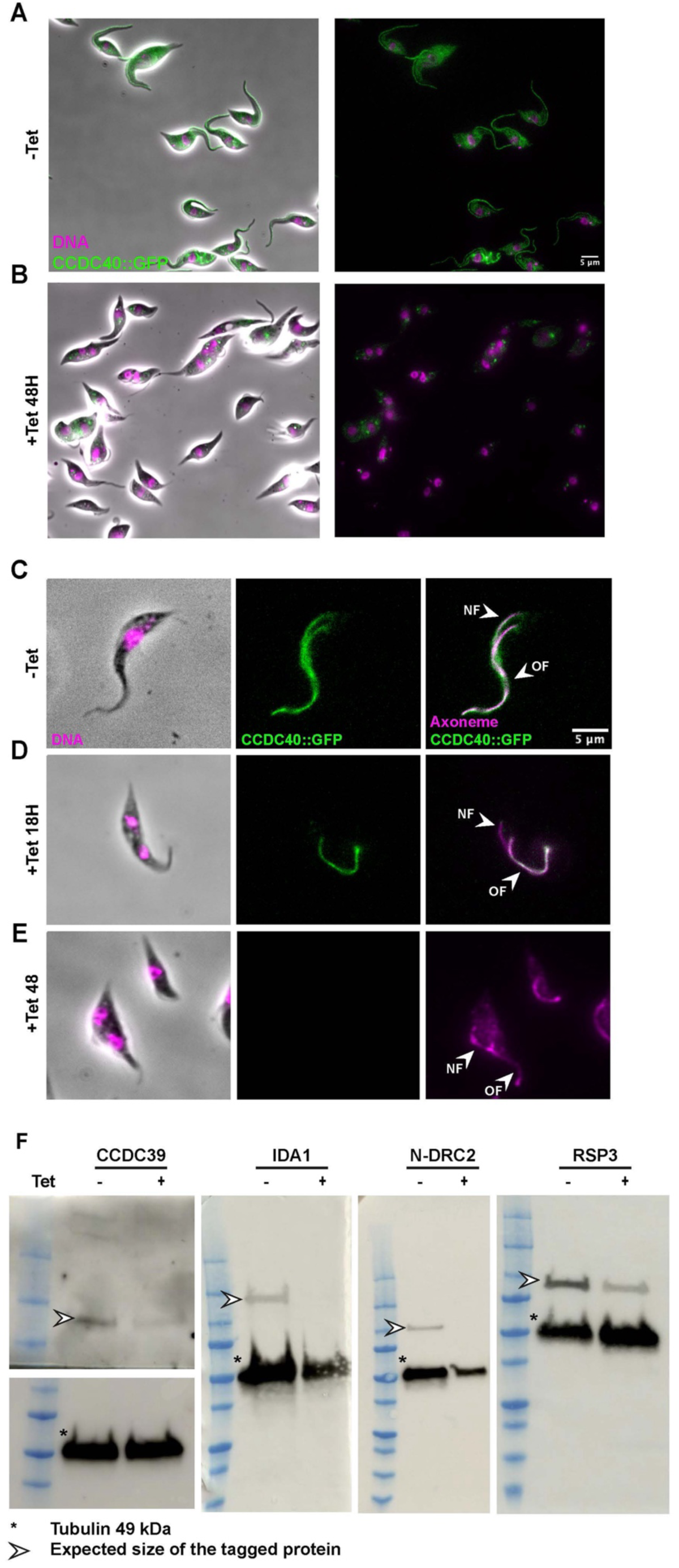
Knockdown of CCDC40 results in its loss from new but not old flagella and impacts associated components. (A-B) Live parasites of the *CCDC40^RNAi1-500^* cells expressing CCDC40::GFP (single-allele tagged, green) were grown without (A) or with (B) tetracycline for 48h, washed, stained with Hoechst (magenta) and observed directly. The fusion protein disappears from flagella upon RNAi knockdown, confirming efficacy of silencing. (C-E) Observation of individual cells of the *CCDC40^RNAi1100-1599^* cell line expressing CCDC40::GFP (single-allele tagged, green) either (C) in non-induced control conditions where both new (NF) and old (OF) flagella are equally positive; (D) after 18h in RNAi conditions where only the new flagellum lacks CCDC40::GFP and (E) after 48h in RNAi conditions when both flagella become negative. Cells were stained with DAPI (DNA, magenta on first panel) and by immunofluorescence with BB2 (detects the Ty1 tags of the CCDC40::GFP fusion protein, green on central and right panels) and mAb25 (axoneme marker, magenta on right panel). (F) Western blot analysis of purified flagella from *CCDC40^RNAi1-500^* cells expressing epitope-tagged versions of either CCDC39::GFP (coiled-coil protein partner of CCDC40), GFP::IDA1 (marker of inner dynein arms), GFP::N-DRC2 (marker of the dynein regulatory complex) and GFP::RSP3 (marker of radial spokes). Cells were grown without (-) or with (+) tetracycline for 72h and flagellar samples were incubated with anti-GFP (CCDC39::GFP, white arrowhead) or BB2 (all the other tagged proteins, white arrowhead) and TAT-1 (tubulin, loading control, indicated by an asterisk).

To visualise the consequences of these disruptions, samples of control and induced *CCDC40^RNAi^* cells were fixed and processed for transmission electron microscopy. Thin sections showed the typical trypanosome flagellum structure in control cells with the 9 doublet microtubules, each carrying outer and inner dynein arms, and the central pair (Fig. 3A). The extra-axonemal paraflagellar rod (PFR) structure is visible, as well as an intraflagellar transport (IFT) train (Fig. 3A). By contrast, the organisation of the flagellum in silenced cells was dramatically affected. Although the 9 doublet microtubules were still present, their regular organisation was perturbed (Fig. 3B). The central pair was still recognisable in some images while the PFR remained present but its structure looked altered (Fig. 3B). These images are compatible with a loss of inter-doublet connection, a phenotype more severe than depletion of DRC components alone (Kabututu et al., 2010). To further explore the nature of the perturbation, non-induced (Fig. 3C,G) and induced (Fig. 3D,H) *CCDC40^RNAi^* cells were expanded by iU-ExM and stained with the polyE anti-tubulin antibody (Casanova et al., 2015). Images were acquired with brighfield microscopy and volumes are presented at Video S1. In non-induced trypanosomes, cross-sections revealed the typical organisation of trypanosome microtubules in both the subpellicular corset and the flagellum, with the 9 doublet microtubules harbouring the typical cylindrical shape (Fig. 3C). Note that PolyE does not detect the central pair (Bonnefoy et al. in preparation), explaining its absence in these images. By contrast, doublet microtubules adopt multiple aberrant independent conformations in induced *CCDC40^RNAi^* cells (Fig. 3D). This messy organisation of the axoneme could reflect broken or interrupted microtubule doublets, something that is difficult to evaluate in cross-sections. Therefore, we examined longitudinal sections by transmission electron microscopy. In contrast to the well organised doublet microtubules of control cells (Fig. 3E), series of doublets exhibiting different orientations were observed in induced *CCDC40^RNAi^*cells (Fig. 3F, arrowheads). Side-views of iU-ExM images brought even nicer resolution since it was possible to follow doublets for several microns. In non-induced samples, doublet continuity is evident (Fig. 3G & Video S1). This was also the case for induced *CCDC40^RNAi^* cells, except that several doublets were frequently distorted and not within the typical cylindrical axis (Fig. 3H, arrowheads & Video S1). We also examined the proximal portion of the axoneme where CCDC40::GFP is absent. Transmission electron microscopy showed a normal organisation for the flagellar pocket, the basal body and the transition zone in both control non induced cells (Fig. S3A) and CCDC40-depleted trypanosomes (Fig. S3B). In conclusion, although axoneme organisation is severely affected with the loss of inner dynein arms, the dynein regulatory complex and the radial spokes, microtubule doublets remain intact but are not connected to each other.

**Figure 3.**
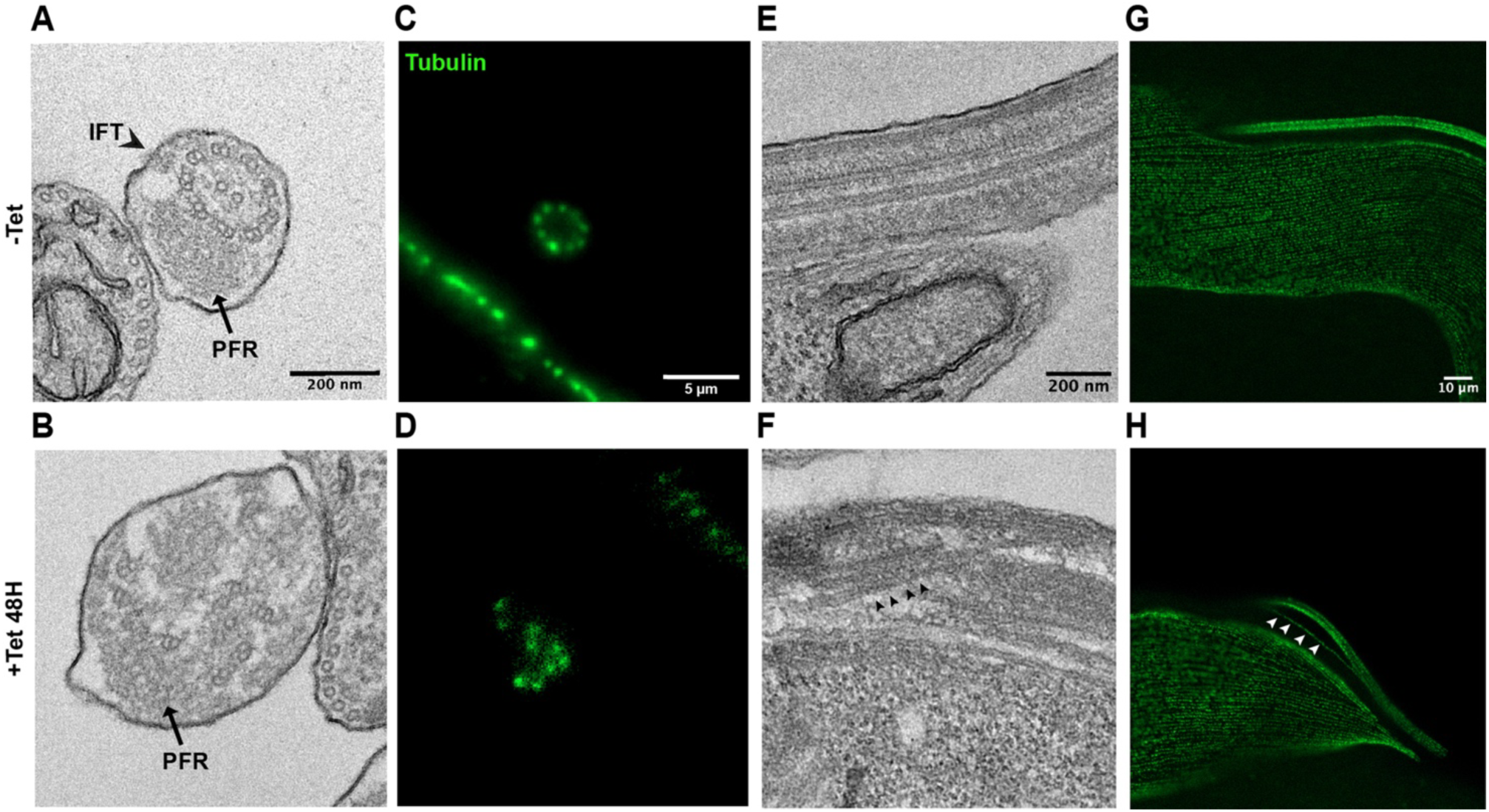
Knockdown of CCDC40 leads to profound alteration in axoneme and associated structures. *CCDC40^RNAi1100-1599^* cell line expressing CCDC40::GFP (single-allele tagged) were grown without (-) or with (+) tetracycline for 48h as indicated. (A-D) Transversal sections of the flagellum viewed by either transmission electron microscopy (A-B) or It-UExM following staining with the polyE antibody that detects tubulin polyglutamylation (C-D). Although the 9 doublet microtubules can be recognised, their organisation is severely disrupted upon silencing. Please note that polyE does not stain the central pair. (E-H) Longitudinal sections of the flagellum viewed by transmission electron microscopy (E-F) or side-views of parasites following It-UExM and staining with the polyE antibody (G-H). While the 9 doublets display conventional organisation in non-induced controls (E,G), they appear disconnected (F,H, arrowheads), albeit neither broken nor interrupted (H) upon CCDC40 silencing.

### Absence of CCDC40 abrogates motility and results in shorter flagella

Looking down the microscope, two obvious phenotypes were observed in induced *CCDC40^RNAi^* trypanosomes: cell paralysis and modified cell shape. First, videos were recorded to monitor cell motility and tracking analysis was performed (Fig. 4). While non-induced trypanosomes showed normal motility (Fig. 4A,B & Video S2, top panels) with the classic combination of individuals displaying persistent, intermediate or tumbling swimming modes (Bargul et al., 2016), *CCDC40^RNAi^* cells induced for 48 hours exhibited little if any motility and mostly drifted with the medium flow (Fig. 4C,D & Video S2, bottom panels). Examination at higher modification showed little or no flagellum beating (Video S2). These results demonstrate that CCDC40 is essential for trypanosome motility.

**Figure 4.**
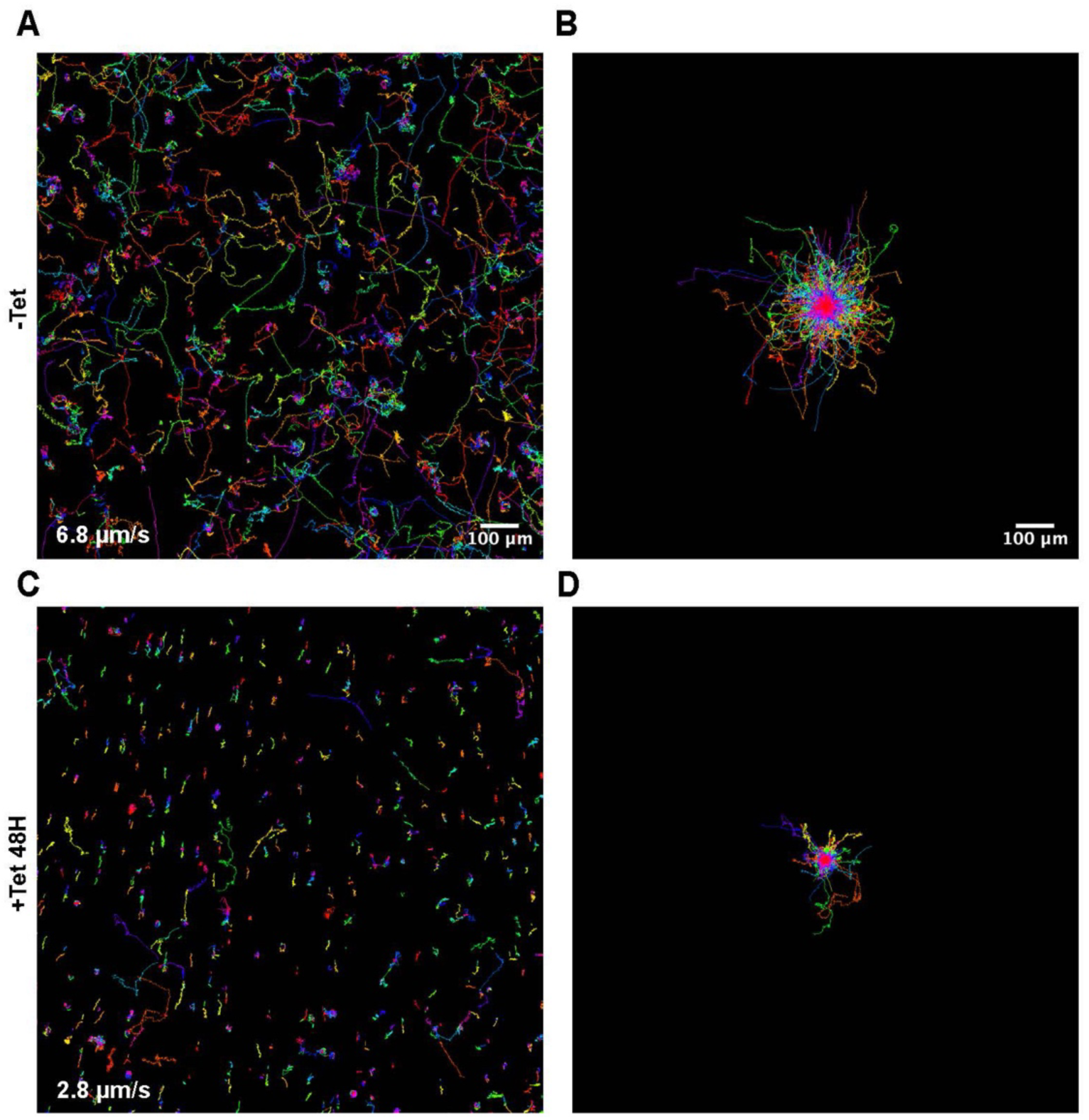
Absence of CCDC40 inhibits trypanosome swimming. (A-D) Trace analysis of a field of *CCDC40^RNAi1-500^* cells imaged at low-magnification. Each trace represents a single trypanosome (colour-coded). While non-induced cells swim actively in all directions (A-B), parasites deprived of CCDC40 drift the medium and rarely display processive motility (C-D). Note that after 48h, a few cells still possess an old flagellum assembled prior to RNAi and swim properly. The average speed is indicated at the bottom left.

Reduced flagellum length was the second striking phenotype observed by light microscopy (Fig. 2B). Non-induced and *CCDC40^RNAi^* cells induced for 48 hours were observed by scanning electron microscopy (Fig. S4). While non-induced controls displayed the classic cellular architecture with the long flagellum attached to the cell body and emerging from the posterior end (Fig. S4A, left panel), a mixture of trypanosomes bearing flagella of different lengths was present after 48h in RNAi conditions, with most of them exhibiting short flagella (Fig. S4B, left panel). Cells growing the new flagellum, connected to its tip to the side of the old one (yellow circles), as well as dividing cells (cleavage furrow indicated with the white arrows) were visible in both conditions (Fig. S4A-B, central panels).

The evolution of flagellum length was monitored during the course of CCDC40 knockdown. Cells were not induced (Fig. 5A) or induced for 18 (Fig. 5B) or 30 (Fig. 5C) hours and up to 6 days (Fig. 5D) followed by staining with the axoneme marker mAb25. The length of the old and of the new flagellum was measured in cells possessing two kinetoplasts and two nuclei (termed 2K2N) that are about to divide since they provide a timely picture of flagellum assembly. In *T. brucei*, the new flagellum grows at a linear rate close to 3 µm/hour while a locking mechanism ensures that the old flagellum cannot incorporate tubulin (Abbuhl et al., 2025; Atkins et al., 2021; Bertiaux et al., 2018b). In control non-induced trypanosomes, elongation of the new flagellum is not yet complete at the time of cell division (Bertiaux et al., 2018b; Robinson et al., 1995), with the new flagellum being ∼20% shorter than the old one (Fig. 5A). After a short induction time (18h), the new flagellum of 2K2N cells looked abnormally short (orange dots) whereas the old one still displayed the normal length (blue dots, Fig. 5B). This is explained by the fact that RNAi degrades mRNAs and that the corresponding proteins disappear according to their own turnover, as shown previously at Fig. 2D. After 30 hours of growth in the presence of tetracycline, two subpopulations were recognised: cells with a long old flagellum and a short new flagellum as seen after 18h (Fig. 5C, top cartoon) and a new category of trypanosomes where both the old and the new flagellum are too short (Fig. 5C, bottom cartoon). These cells correspond to the next generation of those observed after 18h, reflecting trypanosomes that inherited the old or the new flagellum, respectively. Finally, when *CCDC40^RNAi^* cells were grown in knockdown conditions for 6 days, both old and new flagellum were very short, measuring 7 and 6 µm, respectively (Fig. 5D).

**Figure 5.**
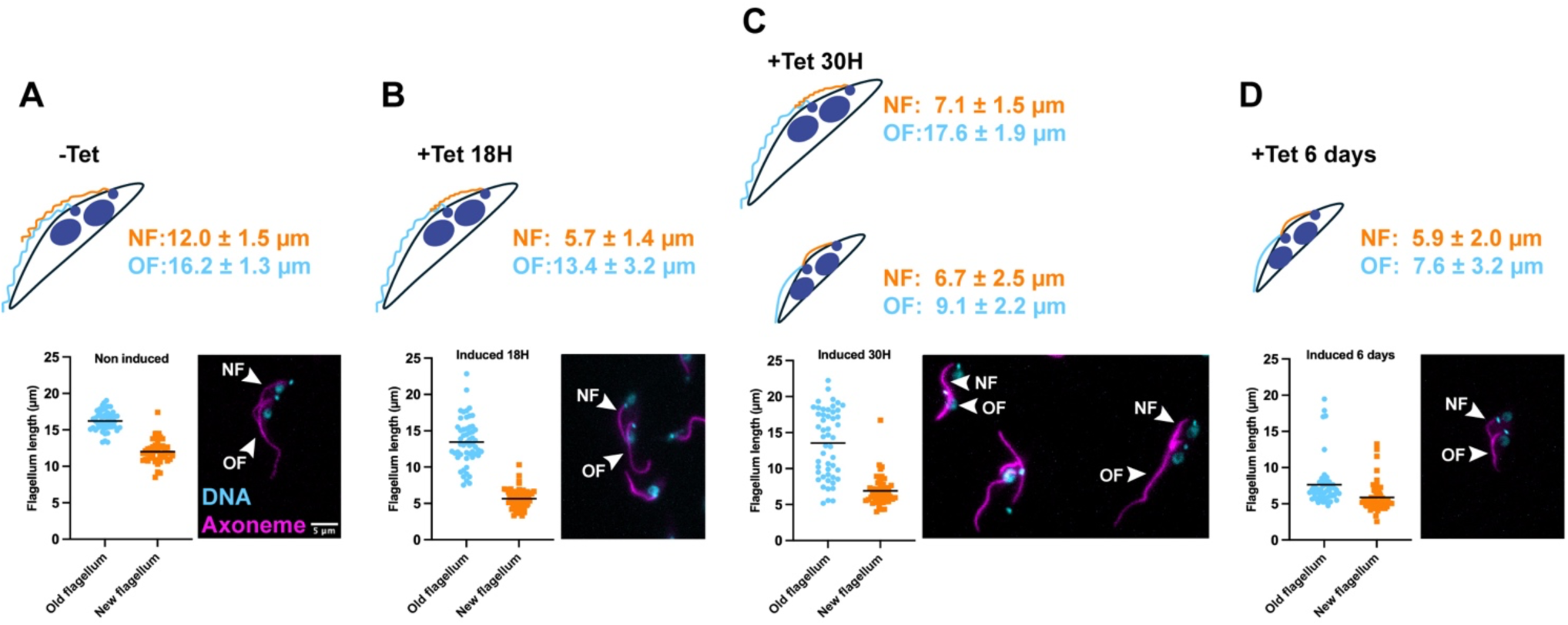
Depletion of CCDC40 impacts flagellum length. (A-D) *CCDC40^RNAi1-500^* cells were grown without (A) or with tetracycline for 18 (B), 30 (C) or 144 (D, 6 days) hours. Cells were fixed and stained with the axoneme marker mAb25 (magenta) and DAPI (blue) to visualise the mitochondrial and nuclear genomes. A total of 50 images of cells at the 2K2N stage (i.e. after mitosis and before cell division) were acquired and the length of both old (OF, blue dots) and new (NF, orange squares) flagella was measured. Cartoons on top of each panel illustrate representative cells for each stage of the induction and the actual average length ± standard deviation of old and new flagella. For the 30h time point, the two groups were defined arbitrarily if the length of their old flagellum was above or below the average flagellum length.

When CCDC40 is absent or mutated, cilia appear shorter in mammalian cells and in *Chlamydomonas* (Becker-Heck et al., 2011; Brody et al., 2025), but this phenotype remains unexplained. Flagellum length in *T. brucei* can be explained with a grow-and-lock model and is amenable to manipulation (Atkins et al., 2021; Bertiaux and Bastin, 2020; Bertiaux et al., 2018b). We therefore set out to understand why flagella are shorter in *CCDC40^RNAi^*cells. First, the rate of tubulin incorporation was measured by expressing a Ty-1 tagged version of alpha-tubulin and monitoring its incorporation in growing flagella. This was achieved by nucleofection of a plasmid expressing tagged alpha-tubulin (Abbuhl et al., 2025) followed by co-staining with antibodies against the epitope tag (green, Fig. 6), the SAXO axonemal marker (Dacheux et al., 2012) and the transition zone marker FTZC (Bringaud et al., 2000)(both antibodies on the same magenta channel, Fig. 6). To avoid signal interference coming from microtubules of the subpellicular corset, these were depolymerised by the addition of 1M NaCl (Robinson et al., 1991) in order to only retain flagella on the slide. Thanks to the presence of the flagella connector, which is also resistant to this treatment (Moreira-Leite et al., 2001), new and old flagella remain tightly associated to each other (Abbuhl et al., 2025). Samples were fixed every two hours and tagged tubulin was indeed incorporated in the new flagellum (Fig. 6C,D). In control cells, the length of the segment labelled increased steadily over time, with an average rate of 1.9 µm per hour (Fig. 6E, green dots), quite close to published values (Abbuhl et al., 2025; Bastin et al., 1999). By contrast, incorporation was much slower in cells depleted of CCDC40 (Fig. 6E, red squares) whose flagella grow at only 0.9 µm per hour, reaching a plateau after 6-8 hours while the control still displayed close to linear growth (Fig. 6E). Hence, the shorter flagellum of induced *CCDC40^RNAi^*cells could be the reflection of a slower growth rate.

**Figure 6.**
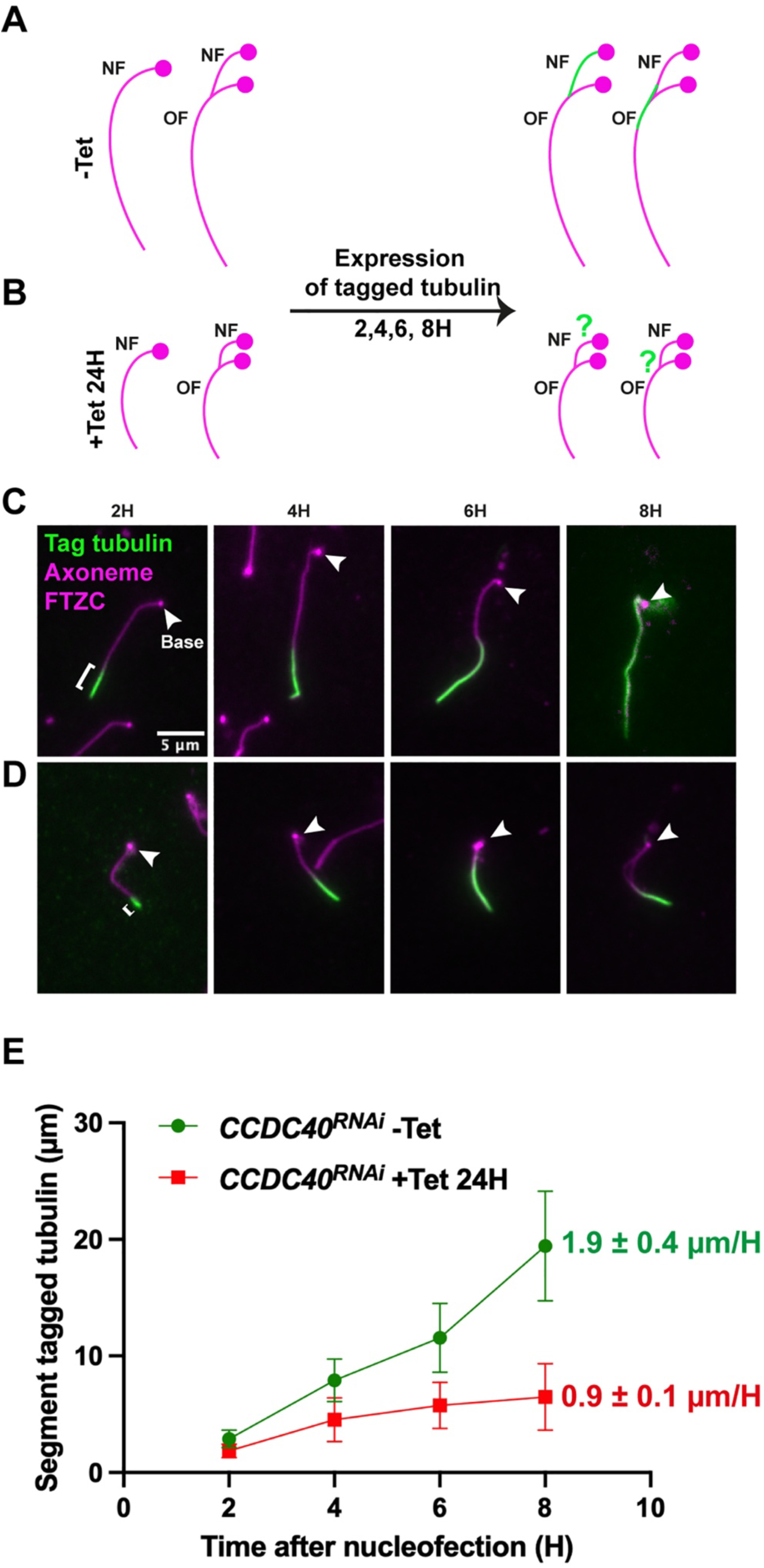
Flagella deprived of CCDC40 show slower incorporation of tubulin. *CCDC40^RNAi1-500^* in pSMOX cell line were grown without (-Tet) or with (+Tet) tetracycline for 24h and nucleofected with a plasmid allowing expression of Ty1-tagged alpha-tubulin. (A,C) The cartoons show flagella (magenta) of cells prior or after expression of tagged tubulin with the expected staining pattern (green) known for control cells (A) (incorporation of newly synthesised tubulin at the distal tip) (Abbuhl et al., 2025). The base of the flagellum is indicated with the magenta spot. Since cells are not synchronised, all phases of the cell cycle can be encountered and tubulin could have been available after (left cartoons) or before (right cartoons) new flagellum growth was initiated. (B,D) To avoid interferences with the cytoplasmic staining of tubulin, flagella were isolated (Abbuhl et al., 2025) and stained with mAb25 (marker of the axoneme, magenta), the anti-FTZC (marker of the base of the flagellum, also in magenta, indicated with the white arrowhead) and BB2 to detect Ty1-tagged tubulin (green) for all the indicated time points. The length of the positive portion is indicated with a white bracket and increases rapidly in non-induced controls (top panels), but more slowly for cells deprived of CCDC40 (bottom panels). (E) Graph representing incorporation of tagged tubulin in growing flagella over time (mean ± standard deviation, n=50). While fast and almost linear growth is visible in non-induced conditions (green spots), incorporation is twice slower in absence of CCDC40 and seems to plateau (red squares).

Since cell cycle progression and flagellum growth are tightly connected, a new flagellum growing too slowly can reach its normal length if cell division is inhibited (Bertiaux et al., 2018b). This can be achieved by incubation in teniposide, a topoisomerase inhibitor that blocks segregation of the mitochondrial genome and hence cell division (Robinson and Gull, 1991). Therefore, non-induced and induced *CCDC40^RNAi^* cells were incubated for 24 hours in teniposide, and flagellum length was measured in cells that had replicated both kinetoplast and nucleus (2K2N) and were about to divide (Fig. 7A-D). In control untreated cells, the length of the new flagellum is about 60% of that of the old one (Fig. 7B, grey bar) but 24 hours after incubation in teniposide, the new flagellum caught up and reacheed the same length as the old one (Fig. 7A-B, majenta bar). Next, *CCDC40^RNAi^* cells induced for 18h were used. At this stage, old flagella still had the normal length but the new one was much shorter at ∼40% of its old counterpart (Fig. 7C-D, grey bar) as observed previously (Fig. 5B). Surprisingly, treatment with teniposide did not modify significantly the length of the new flagellum that remained too short (Fig. 7C-D, majenta bar), suggesting that the slow elongation rate is not the only parameter responsible for the short flagellum phenotype.

**Figure 7.**
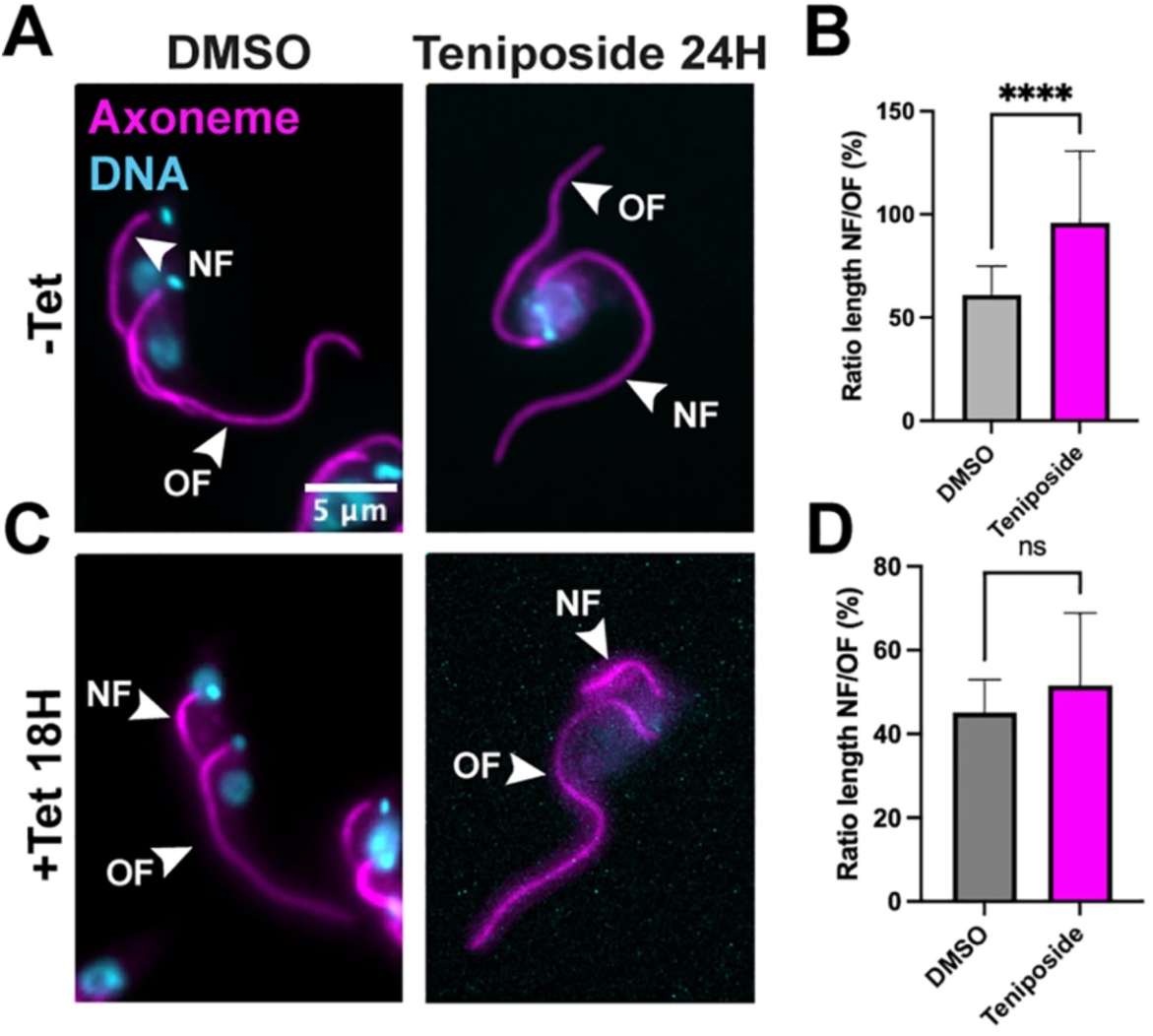
Flagella deprived of CCDC40 cannot elongate further. *CCDC40^RNAi1-500^* trypanosomes grown without (-Tet, A-B) or with (+Tet, C-D) tetracycline for 18h were incubated for an additional 24h with DMSO or 10 mM teniposide, an inhibitor of topoisomerases that blocks kinetoplast segregation and inhibits cell division (Robinson and Gull, 1991). Cells were fixed and stained with the axoneme marker mAb25 (magenta) and DAPI (blue), and the length of old and new flagella (OF and NF, respectively, as indicated with arrowheads) was measured in binucleated cells (n=50). In non-induced and control treated cells, the new flagellum reaches about 60% of the length of the old flagellum (B, grey bar) but inhibition of cell division allows further elongation of the new flagellum, which reaches the length of the old one (B, magenta bar), as expected (Bertiaux et al., 2018b). By contrast, the new flagellum is too short in *CCDC40^RNAi^* induced cells (D, grey bar) and its length is not rescued by blocking cell division (D, magenta bar).

### Possible causes for the short flagellum phenotype

The fact that flagella deprived of CCDC40 could not grow beyond 7-8 µm could be explained by several reasons, and we tried to tackle them one-by-one. IFT is the machinery required for flagellum construction (Prevo et al., 2017) and is a key regulator of flagellum length, as shown upon reduction of IFT train frequency in all species investigated so far (Bertiaux and Bastin, 2020; Bertiaux et al., 2018b; Hoeng et al., 2008; Marshall and Rosenbaum, 2001). Injection of IFT trains takes place at the level of the transition zone at the base of the flagellum (Araujo Alves et al., 2025; van den Hoek et al., 2022). The observed disruption of microtubule doublets (Fig. 3) could affect trafficking and hence reduce protein delivery. *CCDC40^RNAi^* cells were transfected to express an mNG::IFT81 fusion protein as IFT reporter and IFT movement was recorded. Regular IFT trafficking was observed in non-induced cells (Video S3) as expected but also in induced cells (Videos S3), where IFT movement was clearly visible, even on disrupted microtubules (Video S3). Therefore, a reduction in IFT trafficking cannot explain the short flagellum phenotype.

The *T. brucei* flagellum displays heterogeneities along its length, with a proximal and a distal segment of different composition for the docking complex of outer dynein arms (Bonnefoy et al., 2024; Bonnefoy et al., 2018; Edwards et al., 2018; Fort et al., 2025). The short flagellum phenotype could reflect the absence of one or the other segment. We therefore transformed the *CCDC40^RNAi^* cell line to produce tagged versions of pDC1 and dDC1 for the proximal and distal docking complex, respectively. In control non-induced cells, growing flagella displayed the expected combination of proximal and distal complex components as early as they emerge from the flagellar pocket, keeping the same proportions during construction (Fig. S5A-B) as reported previously (Edwards et al., 2018). CCDC40 knockdown cells possessed shorter flagella but the distribution of proximal and distal elements kept the right proportions (Fig. S4C-D). Therefore, the phenotype is not due to the absence/reduction of one or the other proximal/distal segments which are simply shorter altogether.

Flagella mature over at least one cell cycle in trypanosomes (Bertiaux and Bastin, 2020; Farr and Gull, 2009). We used a polyclonal antibody to examine the location of the maturation marker FLAM8 (Calvo-Alvarez et al., 2021; Subota et al., 2014). This protein is found at the tip of axoneme microtubules in low concentrations in short flagella and its amount increases as they elongate. At cell division, the amount of FLAM8 at the distal end of new flagella was about 65% compared to that at the old flagellum (Fig. 8A, C, green). By contrast, FLAM8 already reached the maximum concentration equivalent to the old flagellum in induced *CCDC40^RNAi^* cells (Fig. 8B, C), suggesting that this flagellum has already matured, potentially explaining why it does not grow further.

**Figure 8.**
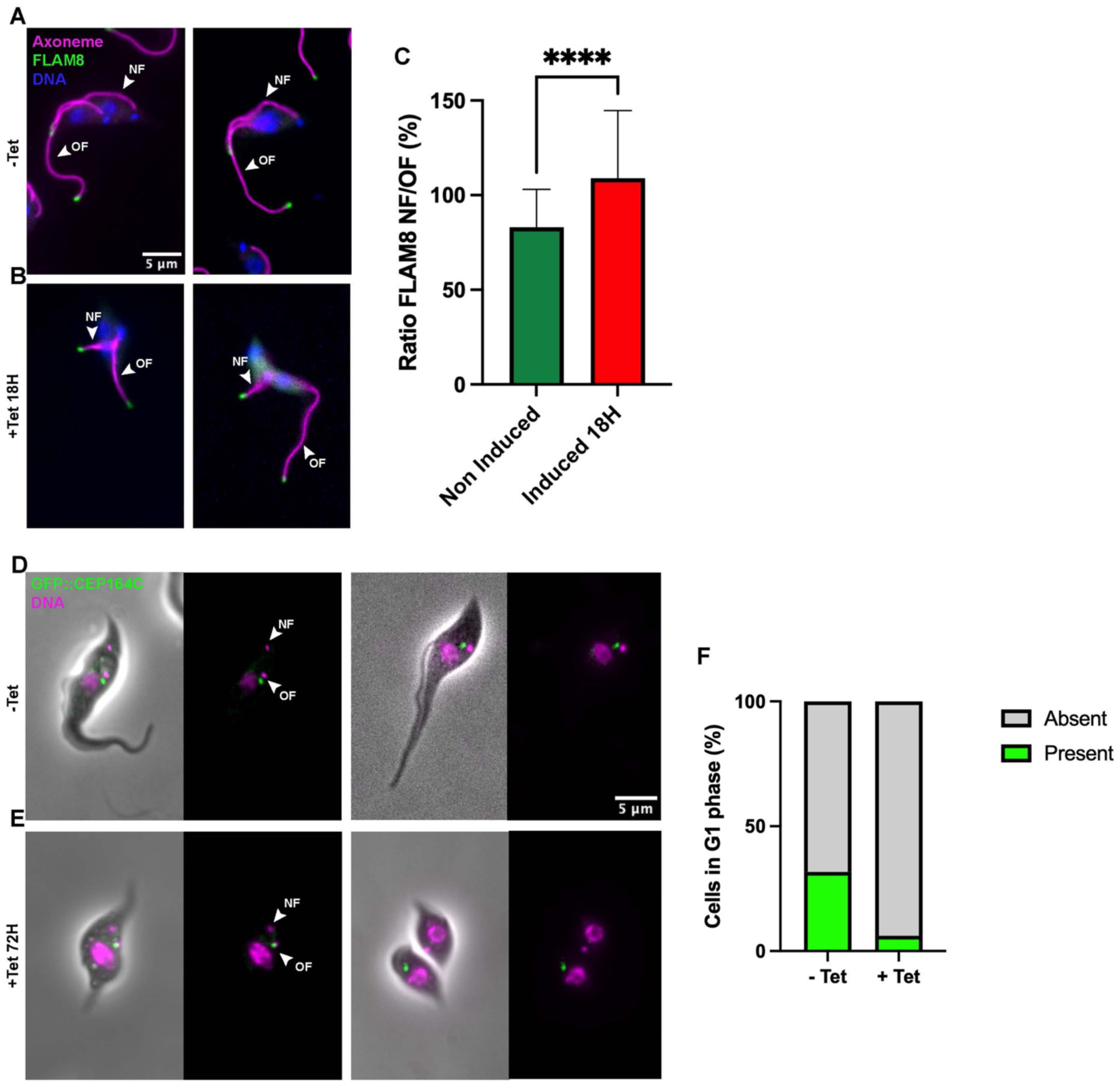
Flagella deprived of CCDC40 acquire the locking and maturation proteins. (A-B) *CCDC40^RNAi1-500^* trypanosomes grown without (-Tet, A) or with (+Tet, B) tetracycline for 18h were fixed and stained with the axoneme marker mAb25 (magenta), an antibody against the tip protein FLAM8 (green) and DAPI (blue). Two representative biflagellated cells are shown for each condition. The intensity of the FLAM8 signal was measured at the tip of each flagellum of binucleated cells and the ratio of these intensities between new and old flagella is presented at panel C (mean ± standard deviation, n=50). While the amount of FLAM8 is always lower at the tip of the new flagellum versus the old one in control cells (C, green bar) as expected (Bertiaux et al., 2018b), it is found equally on both flagella in induced cells (C, red bar). (D-F) *CCDC40^RNAi1-500^* trypanosomes expressing the locking protein GFP::CEP164C were grown without (-Tet, D) or with (+Tet, E) tetracycline for 72h, washed, stained with Hoechst (magenta) and observed directly. Two representative images are shown for each condition. The fusion protein GFP::CEP164C (green) is present at the base of the old flagellum (OF) and not the new flagellum (NF) in all biflagellated cells in both conditions (D,E). However, its removal following cell division is much faster in the absence of CCDC40 (F, percentage of G1 cells with mNG::CEP164C staining; for non-induced, n=467 and for induced, n=309).

During new flagellum assembly, the old flagellum is locked to prevent untimely elongation (Bertiaux et al., 2018b). This is controlled by the CEP164C protein that associates firmly to the transition fibres of the basal body of exclusively the old flagellum (Atkins et al., 2021). The early acquisition of the FLAM8 maturation marker might reflect a premature locking event of the new flagellum that would take place before cell division and therefore before normal length is reached. This could also explain the plateau in tubulin incorporation observed in the new flagellum of *CCDC40^RNAi^* cells (Fig. 6). CEP164 fused to GFP was therefore expressed endogenously in the *CCDC40^RNAi^* cell line and its localisation was monitored. CEP164C was present in all biflagellated cells but exclusively associated to the base of the old flagellum in both control (Fig. 8D) and CCDC40-depleted cells (Fig. 8E). CEP164C does not remain associated to the old flagellum for ever and is removed after cell division, probably to allow growth in the G1 phase to compensate for the slight shrinking observed during the locking period (Abbuhl et al., 2025; Abeywickrema et al., 2019; Atkins et al., 2021). The protein was correctly removed in G1 cells and even faster in the case of induced cells (Fig. 8F). We conclude that a premature locking is not the cause of the short flagellum phenotype in the absence of CCDC40.

### Immotile cells with short flagella divide and grow normally

We have showed that depletion of CCDC40 leads to the assembly of short flagella and consequently to the formation of shorter cells. The internal flagellar organisation is severely disrupted, and cells fail to swim (Fig 4). Cell growth was followed for more than a week, and surprisingly non-induced and induced parasites grew at exactly the same rate (Fig. 9A), revealing that *CCDC40^RNAi^* cells do not require motility for cell division. This goes against the dogma that motility is essential for trypanosome cytokinesis but could be explained by the shorter size of the cell body and the simpler path needed to separate the future daughter cells. Knowing the role of the FAZ in cell division (Kohl et al., 2003; Zhou et al., 2011), its presence and distribution was examined in non-induced and induced *CCDC40^RNAi^* cells using the marker antibody L3B2 that recognises FAZ1, a major FAZ component (Kohl et al., 1999; Vaughan et al., 2008). In control cells, FAZ1 was found all along the flagellum on the cell body side, starting at the flagellum exit point from the flagellar pocket and running until the anterior end of the cell body (Fig. 9B) as expected. This location was conserved in *CCDC40^RNAi^*cells induced for 2 days, the only difference being a shorter FAZ associated to the shorter flagellum (Fig. 9C). This demonstrates that the FAZ is still assembled and follows the flagellum, hence is able to perform its guide function for cytokinesis normally.

**Figure 9.**
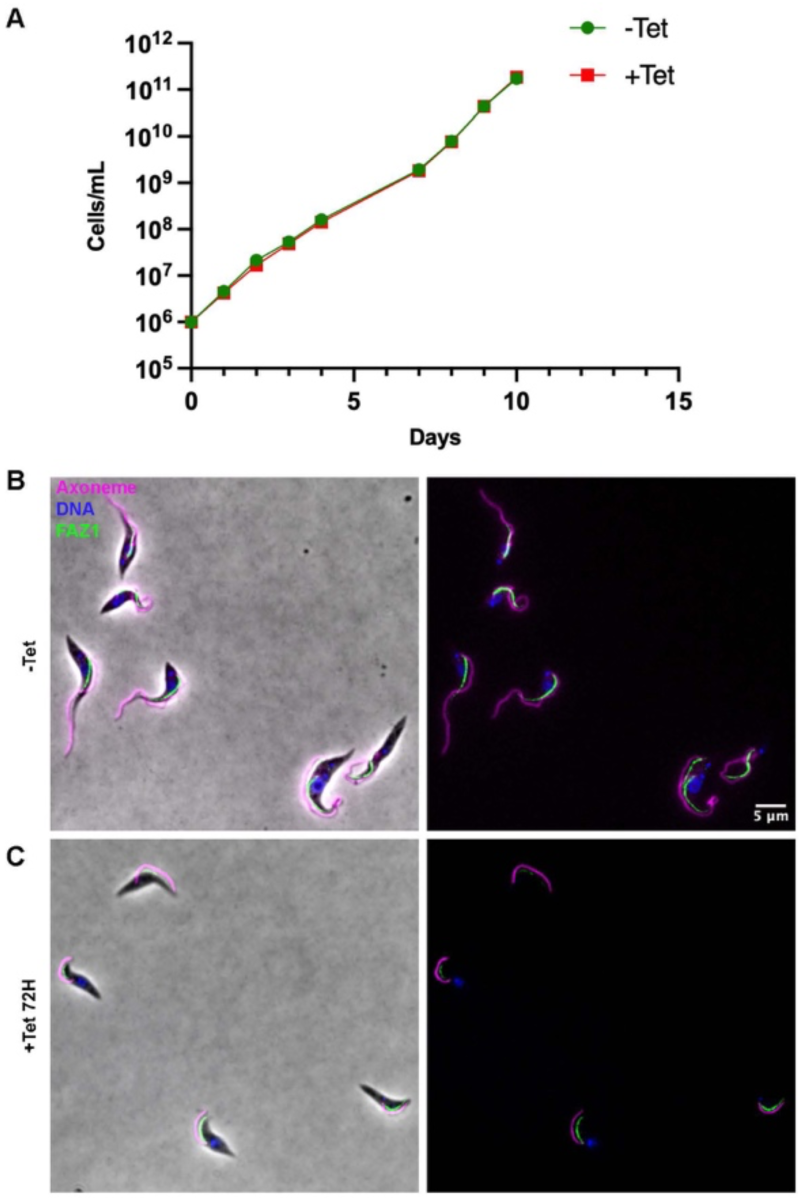
Despite being shorter and unable to swim, CCDC40 depleted trypanosomes grow normally and assemble a FAZ filament. (A) Cumulative growth curve of *CCDC40^RNAi1-500^* cells grown in absence (green) or presence (red) of tetracycline for 10 days. Non-induced and induced parasites display the same growth rate. (B,C) *CCDC40^RNAi1-500^* cells were grown without (B) or with (C) tetracycline and stained with the axoneme marker mAb25 (magenta), the FAZ marker FAZ1 (green) and DAPI. The FAZ filament follows the flagellum on the cell body side and ends at the anterior part of the cell in control (top panels) and RNAi conditions (bottom panels), despite the shorter flagellum length.

## Discussion

Cryo-electron microscopy analyses of splayed axonemes from *Chlamydomonas* and human ciliated organoids (Walton et al., 2023) as well as from *T. brucei* (Xia et al., 2025) revealed that CCDC39 and CCDC40 form a dimer stretched out at the surface of the A-tubule that serves as a platform for attachement of inner dynein arms, radial spokes and the dynein regulatory complex. The actual length of the dimer is responsible for the regular spacing of the 96-nm repeats (Oda et al., 2014). Following tagging of both alleles of the trypanosome *CCDC40* gene, iUExM revealed nicely the repetitive pattern of the fusion protein along the axoneme, with a periodicity matching the structural prediction. This result shows the power of iUExM to analyse the molecular composition of cytoskeletal components without requiring sophisticated microscopy technologies (Louvel et al., 2023). Double tagged cells grow and behave normally, formally demonstrating that the fusion protein containing GFP and 6 copies of the Ty1 tag is functional.

RNAi knockdown of CCDC40 has a profound impact on axoneme organisation with depletion of numerous components, resulting in the loss of connection between microtubule doublets. In *T. brucei*, individual knockdown of the DRC (Hutchings et al., 2002; Kabututu et al., 2010), the inner dynein arms (Springer et al., 2011) or the radial spokes (Ralston et al., 2006; Ramakrishnan et al., 2023) does not produce such a strong phenotype, suggesting that it is the combined loss of these elements that is responsible for the severe axoneme phenotype reported here. These results show that CCDC40 distribution and function is conserved in trypanosomes, further validating them as amenable models to investigate ciliary genes mutated in human patients (Bonnefoy et al., 2018; Coutton et al., 2018; Dacheux et al., 2023).

*CCDC40^RNAi^* cells exhibit a much shorter flagellum, a feature previously reported in *Chlamydomonas* mutants and in human patients, both for motile cilia in epithelial cells and for the flagellum of spermatozoa (Aprea et al., 2023; Brody et al., 2025). However, the origin of this defect is unclear. In trypanosomes, depletion of central pair proteins, dynein arms or DRC subunits impacted strongly on flagellum beating and cell motility, but had limited or no impact on flagellum length (Absalon et al., 2007; Kabututu et al., 2010; Ralston et al., 2006; Springer et al., 2011). By monitoring tubulin incorporation, we have shown that the flagellum growth rate is 2-3 fold slower in *CCDC40^RNAi^* cells, reaching a plateau prematurely (Fig. 6). A slower flagellum growth rate was also observed upon knockdown of the IFT kinesins, resulting from a reduced frequency of IFT train injection. This phenotype was compensated by inhibiting cell division with teniposide allowing further flagellum elongation (Bertiaux et al., 2018b), which was not the case here. The shorter flagellum length is not due to premature locking, which could have halted flagellum elongation, since the locking protein CEP164C (Atkins et al., 2021) is detected exclusively at the base of the old flagellum in biflagellated cells, exactly as in controls. Nevertheless, CEP164C is removed much faster after cell division in *CCDC40^RNAi^* cells. The distal tip protein FLAM8 is considered as a marker of flagellum maturation since it accumulates during assembly and reaches maximal concentration when the new flagellum has reached his final length (Calvo-Alvarez et al., 2021; Subota et al., 2014). By contrast, *CCDC40^RNAi^* cells acquire an equivalent amount of FLAM8 at the tip of both flagella at the time of cell division. This could be an explanation for the limited length of the flagellum, but one has to keep in mind that absence of FLAM8 does not affect flagellum length (Calvo-Alvarez et al., 2021; Subota et al., 2014). The faster acquisition of FLAM8 in the new flagellum as well as the accelerated removal of CEP164C from the base of the old flagellum suggest that these processes could be related to flagellum length. If FLAM8 reaches the distal tip by diffusion as the EB1 tip protein does in *Chlamydomonas* (Harris et al., 2016), it would do so faster in a shorter flagellum compared to a normal one, possibly explaining why it reaches maximum concentration earlier since cell cycle progression is unchanged. Similarly, if CEP164C removal from the base of the flagellum relies on a process related to protein diffusion in that same flagellum, a shorter distance would allow a faster removal. Elucidating the mechanisms controlling addition or removal of proteins will be the subject of promising future research.

The trypomastigote shape of bloodstream and procyclic trypanosome stages is characterised by the emergence of the flagellum from the posterior end of the cell (Hoare, 1966). From there, the flagellum is attached to the cell body, a configuration presumably evolved for optimal swimming in viscous environments such as those encountered in blood and tissues of mammalian hosts (Bargul et al., 2016; Heddergott et al., 2012), or the midgut of the tsetse fly (Rotureau et al., 2011). It also favours rapid capture of surface-bound antibodies and clearance via the flagellar pocket (Engstler et al., 2007). Coherently, disruption of attachment leads to drastic drops in swimming directionality despite the presence of an actively beating flagellum (Rotureau et al., 2014; Woods et al., 2013).

This configuration involves tight constraints on cytokinesis since the division plane is initiated close to the tip of the new flagellum and needs to bisect the cell from anterior to posterior for a long distance (Wheeler et al., 2019)(Fig. 10A). As described in the introduction, the flagellum contributes to this process by several ways, each of them being essential. Inhibition of flagellum motility blocks cytokinesis, but without preventing reentry in the cell cycle, followed by S phase and mitosis, resulting in the formation of large multinucleated cells (Fig. 10B) (Branche et al., 2006; Broadhead et al., 2006; Kabututu et al., 2010; Ralston et al., 2006; Springer et al., 2011)). The new flagellum functions acts as a guide for elongation of the FAZ filament, which in turn positions the cell division axis (Zhou et al., 2011). When flagella are not assembled upon silencing of IFT components, the FAZ does not elongate and short cells are formed, which also fail to divide (Kohl et al., 2003) (Fig. 10C). By contrast, reducing FAZ length shifts the flagellum in an anterior position relative to the nucleus but it keeps on beating, allowing normal cell division (Hayes et al., 2014)(Fig. 10D).

**Figure 10.**
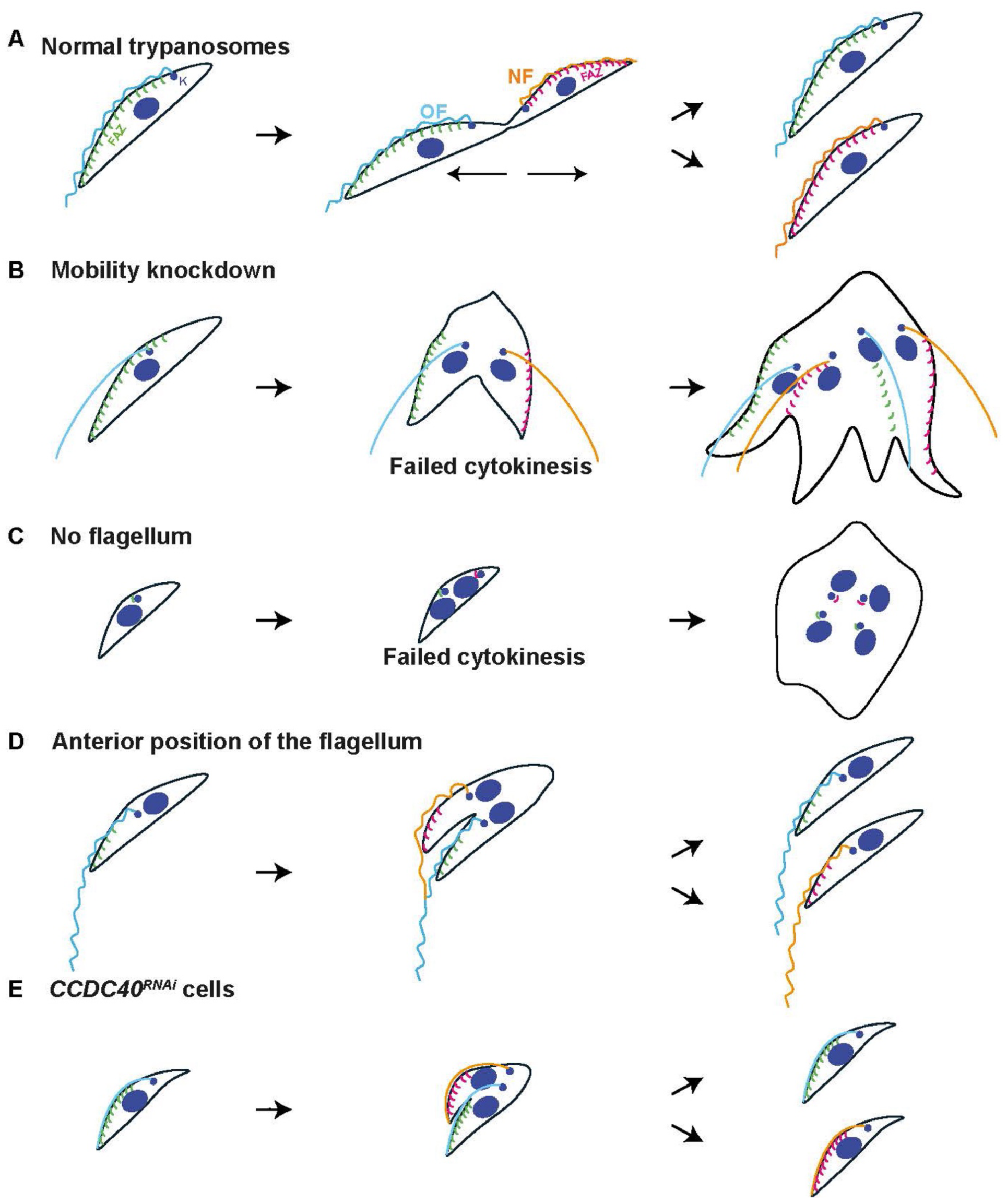
Model explaining the involvement of flagellum presence and motility in trypanosome division. (A) In normal trypanosomes, the new flagellum is attached along most of the cell body via the FAZ and cell motility is required to complete cytokinesis and produce two daughter cells (Bertiaux et al., 2018b). (B) When flagellum beating is inhibited, the kinetoplast is positioned closed to the the nucleus and cytokinesis fails. Nevertheless, basal bodies, kinetoplasts and nuclei keep on replicating, leading to the emergence of multinucleated and multiflagellated cells with a large number of kinetoplasts (Branche et al., 2006; Broadhead et al., 2006; Ralston et al., 2006). (C) When flagellum assembly is blocked, basal bodies still duplicate but the FAZ does not elongate. Cells do not divide, yet duplicate the kinetoplasts and undergo mitosis (Kohl et al., 2003). (D) When the flagellum is in an anterior position relative to the nucleus, a short FAZ is assembled and flagellum beating remains active, allowing normal cell division (Hayes et al., 2014). (E) In *CCDC40^RNAi^* cells, flagella are three-times shorter than normal but remain attached to a shorter cell body via a short FAZ. Although flagella are immotile, these “mini-cells” managed to divide at a normal rate.

The unexpected normal growth of paralysed *CCDC40^RNAi^*cells could be explained by the short cell body length resulting from the short flagellum phenotype (Figure 10E). Since cytokinesis in *CCDC40^RNAi^* cells needs to process through a much shorter distance, active flagellum beating might not be required for its completion. The situation can be compared to the early stages of IFT knockdown where shorter flagella are constructed, in association with a shorter FAZ, but these cells can still divide (Bertiaux et al., 2018b; Kohl et al., 2003). Another parallelism can be drawn with the related Kinetoplastid parasite *Leishmania*, where the flagellum is only attached to a short portion of the cell body within the flagellar pocket neck region and is free for the rest of its length. Inhibition of flagellum assembly is accompanied by a change in cell body shape that looks stumpier but this does not impact neither cell division nor cell growth (Adhiambo et al., 2005; Beneke et al., 2019; Sunter et al., 2018). This is coherent with the simpler architecture of the cell body and the straightforward path for the cell division axis (Hair et al., 2024; Wheeler et al., 2019).

Another hypothesis for the role of the flagellum in cell division posits that axoneme anatomy should be intact to allow proceeding to the next cell cycle (Ralston et al., 2011). This was based on the expression of point mutations of the dynein light chain 1 (LC1), which interferes will flagellum beating without visibly affecting the structural organisation of dynein arms on the axoneme. Since these cells were able to complete cell division even at the bloodstream stage, it was proposed that the cytokinesis defects reported in multiple knockdowns were a consequence of axoneme perturbation rather than of reduced motility (Ralston et al., 2011). This explanation is not compatible with *CCDC40^RNAi^* cells that divide normally despite the dramatic modifications of axoneme organisation.

Altogether, our investigations of CCDC40 in *T. brucei* confirm its conserved architecture and function across eukaryotes and further validate trypanosomes as useful models to study cilia and ciliary proteins. They provide insights as to why cilia and flagella are shorter in the absence of CCDC40 and reveal that the flagellum motility requirement for trypanosome cytokinesis (in contrast to *Leishmania*) is probably explained by the intricate configuration of the trypomastigote stage where cell division has to progress through a very long cell body.

## Material and methods

### Cell culture

Procyclic trypanosome cell lines expressing the T7 RNA polymerase and the tetracycline-repressor either with two markers (29-13) (Wirtz et al., 1999) or a single marker (pSMOX) (Poon et al., 2012) were cultured at 27° in SDM-79 medium (Brun and Schonenberger, 1979) at pH 7.4, complemented with 10% fetal bovine serum and Hemin (7.5 mg/l). For growth curves, cells were counted every 24h with a Luminex® Guava® Muse® flow cytometer and then diluted to 1 x 10^6^ cells/ml.

### Vectors and transfection

The *CCDC40* gene (Tb927.10.11250) was tagged at its 3’ end with the endogenous PCR strategy using pPOTv6GFP with a puromycin resistance marker and the second allele was tagged with the pPOTv6mNG with a nourseothricin resistance cassette (Dean et al., 2015). These vectors contain the coding sequence for GFP or mNG with 6 copies of the Ty1 epitope tag (Billington et al., 2023; Dean et al., 2015)(Fig. S2). For RNAi, two different fragments of the *CCDC40* gene were generated by chemical synthesis by GeneCust Europe (Dudelange, Luxembourg), the first one from nucleotide 1-500 of the coding sequence (*CCDC40^RNAi1-500^*) and the second one from 1100 to 1599 (*CCDC40^RNAi1100-1599^*). Genecust cloned these fragments into the pZJM vector (Wang et al., 2000), allowing tetracycline-inducible expression of dsRNA generating RNAi upon transfection in the 29-13 (Wirtz et al., 1999) or in the pSMOX (Poon et al., 2012) recipient cell lines. The dsRNAs are expressed from two tetracycline-inducible T7 promoters facing each other in the pZJM vector. Sequences were selected using the RNAit algorithm to ensure that the fragment lacked significant identity to other genes to avoid cross-RNAi (Redmond et al., 2003). Prior to transfection the plasmid was linearized with NotI. For transfection, 1 x 10^8^ cells were harvested and then transfected with 10 µg of linearized plasmid using an Amaxa nucleofector II (Burkard et al., 2007). Transfectants were grown in media with the appropriate antibiotic concentration. Since the same phenotype was observed with both RNAi segments, the term *CCDC40^RNAi^* was used throughout the text for the sake of lisibility but the actual name of the cell line used is given in each figure legend.

To express mNG::IFT81 and mNG::dDC1, the *CCDC40^RNAi^* cell line was transformed with the p2675mNGIFT81 plasmid linearized with the enzyme XcmI (Bertiaux et al., 2018a) and with the p2675mNGdDC1 plasmid linearized with the enzyme ClaI (Bonnefoy et al., 2024), respectively. To express pDC1, the long primer strategy was used with plasmid p2675dDC1 as template. All the other tagging experiments were performed with the same PCR strategy using either pPOTv6 or pPOTv7 as template (Dean et al., 2015). These vectors contain the coding sequence for GFP with 6 copies of the Ty1 epitope tag (pPOTv6) or only GFP (pPOTv7) and a puromycin or nourseothricin resistance cassette (Billington et al., 2023; Dean et al., 2015). For CCDC39 (coiled-coil domain containg protein 39), pPOTv7GFP was used while pPOTv6GFP was used for IDA1 (inner dynein arm 1), N-DRC2 (nexin-dynein regulatory complex 2), RSP3 (radial spoke protein 3) and CEP164C. The sequence of primers is presented at Table S1. Cells were nucleofected with the PCR products using an Amaxa nucleofector II as above (Burkard et al., 2007). Transfectants were grown in media with the appropriate antibiotic concentration.

### Live imaging

Trypanosomes were washed three times with phosphate-buffered saline, Hoechst (1 µg/mL) was included in the first wash to stain the DNA. Cells were spread on slides and imaged immediately (Billington et al., 2023). Acquisition was performed with a Photometrics PRIME 95B^TM^ back illuminated scientific CMOS camera using a Leica DMI4000 microscope (100x objective, NA=1.4) and analysed with ImageJ (Schindelin et al., 2012).

### Immunofluorescence assays

Trypanosomes were harvested at densities between 5 x 10^6^ – 1 x 10^7^ cells/mL. For whole cell analysis, cells were transferred onto poly-L-lysine coated slides, left in humid chamber for 10 minutes and then fixed with PFA4% for 5 min and methanol at −20°C for 5 min. To extract cytoskeletons, cells were settled on poly-L-lysine slides for 10 min, then treated with 0.5% Nonidet P-40 in PEM buffer (2 mM EGTA; 1mM MgSO_4_; 0.1 M, pH6.9, PIPES) for 5 minutes and fixed with methanol at −20°C for 5 minutes (Robinson et al., 1991). Extraction of flagella was achieved by treating detergent extracted cytoskeletons with 1M NaCl twice for 5 – 10 minutes before fixation. Post fixation samples were rehydrated in PBS for 15 minutes. Primary and secondary antibodies (Table S1) were diluted in 0.1%BSA in PBS and incubation last 60 min in humid chamber. DNA was stained with 4′,6-diamidino-2-phenylindole (DAPI, 10 µg/ml in PBS) or Hoechst for 5 minutes. Glass cover slips were mounted with ProLong^TM^ Gold antifade reagent. Images were acquired as described above.

### Western blot analysis

Procyclic cells were harvested at densities between 2 x 10^7^ cells/mL. To obtain whole cell lysates, cells were resuspended in 200 µl hot Laemmli buffer and then boiled at 100°C for 5min. For flagella fraction the cell pellet was treated with 1% NP-40 in PEM buffer containing protease inhibitor for 2 minutes. The pellet was harvested by centrifugation and then resuspended in 1M NaCl for 2 minutes. Then the pellet was harvested by centrifugation and boiled in Laemmli buffer for 5min at 100°C. Samples were separated by sodium-dodecyl sulfate-polyacrylamide gel electrophoresis (SDS-PAGE) in Mini-PROTEAN TGX Precast gels from BIO-RAD with the BIO-RAD running system in Tris / Glycine / SDS – buffer. Proteins were then transferred onto nitrocellulose membranes using the Trans-Blot Turbo transfer system of BIO-RAD. Membranes were blocked at room temperature in 5% milk in PBS for 1 hour. Primary antibodies (Table S1) diluted in 2.5% powder milk in PBS were incubated overnight at 4°C. Thereafter membranes were washed 3x in 0.1% PBS-Tween. Secondary antibodies were added in PBS 60 minutes at room temperature. Membranes were then washed in 0.1% Tween in PBS 3 times for 10 minutes. The blots were then incubated with ECL Select^TM^ Western Blotting Detection Reagent for 5 minutes and revealed with an Amersham^TM^ ImageQuant 800. Signal intensity was estimated with the ImageQuantTL® software.

### Motility assay

Cells were cultured at a low density of approximately 1 × 10⁶ cells/mL, and 5 µL of the suspension was deposited onto a glass slide within a Gene Frame. A coverslip was placed on top of the Gene Frame to create a chamber, allowing the parasites to swim freely. Images were acquired using a 10x objective on a Leica DMI4000 microscope, with a 2 ms exposure time. A total of 512 frames were captured at 200 ms intervals. Data were analyzed using the TrackMate plugin in ImageJ (Tinevez et al., 2017; Wheeler, 2017).

### Transmission electron microscopy

Procyclic cells were fixed overnight at 4°C in a solution containing 2.5% glutaraldehyde and 2% formaldehyde in 0.1 M PHEM (PIPES, HEPES, EGTA, MgSO4) buffer. After fixation, cells were washed three times with 0.1 M PHEM buffer 5 min each and post-fixed with 1% osmium and 1.5% potassium ferrocyanide in 0.1 M PHEM buffer for 1 h. Samples were treated for 30 min with 1% tannic acid and 1 h with 1% osmium tetroxide, rinsed in water and incubated in 1% uranyl acetate dissolved in 25% ethanol for 30 min. Cells were dehydrated in an ethanol series comprising 25%, 50%, 75% and 95% ethanol solutions (5 min each), followed by three immersions in 100% ethanol (10 min each). Cells were embedded in epoxy resin after 48 h at 60 °C of polymerization. 50-70 nm sections were obtained using an EM UC6 ultra-microtome (Leica) and stained with 2% uranyl acetate in milli-Q water and 80 mM lead citrate in milli-Q water before imaging. The sample was imaged using a Tecnai BioTWIN 120 microscope (FEI) equipped with a MegaView II camera (Arecont Vision).

### Iterative ultrastructure expansion microscopy (iU-ExM)

For iU-ExM, 10^7^ *T. brucei* cells were washed twice in PBS and were adhered to poly-L-lysine-covered 12-mm coverslips for 15 minutes. The next steps were done exactly as described (Louvel et al., 2023). Acquisition was performed either with a Dual camera sCMOS Hamamatsu ORCA FLASH 4.0 v2.0 using a Widefield Zeiss microscope (63x objective) or with a Photometrics PRIME 95BTM back illuminated CMOS camera using a Leica DMI4000 microscope (63x objective) and analysed with ImageJ (Schindelin et al., 2012).

### Scanning electron microscopy

For scanning electron microscopy, samples were fixed overnight at 4°C with 2.5% glutaraldehyde in 0.1 M cacodylate buffer (pH 7.2) and post-fixed in 1% OsO4 in 0.1 M cacodylate buffer (pH 7.2)(Bertiaux et al., 2018b). After serial dehydration, samples were critical-point dried (Emitech K850 or Balzers Union CPD30) and coated with gold (Jeol JFC-1200 or Gatan Ion Beam Coater 681). Acquisitions were made in a JEOL 7600F microscope.

### Inibition of cell division

For inhibition of cell division, teniposide, a topoisomerase II inhibitor was dissolved in DMSO and added to trypanosome cultures at a final concentration of 200 μM during 24 hours (Bertiaux et al., 2018b; Robinson and Gull, 1991). In the control flask, the same volume of DMSO was added (68 μL).

### Tubulin incorporation

To monitor tubulin incorporation in growing flagella, *CCDC40^RNAi^* cells grown in absence or presence of tetracycline were nucleofected with 10 µg of a circular version of the plasmid pHD4301xTy1-tubulin allowing expression of tagged alpha-tubulin (Abbuhl et al., 2025). Cells were returned to the culture medium and grown for 2, 4, 6 and 8h in regular conditions. At the end of each incubation time, cells were washed and flagella were extracted as published (Abbuhl et al., 2025). They were stained with the axoneme marker mAb25 (Dacheux et al., 2012), the antibody against the flagellar transition zone component FTZC (Bringaud et al., 2000), the BB2 antibody to detect the Ty-1 epitope (Bastin et al., 1996) present in alpha-tubulin and DAPI before imaging with the same Leica DMI4000 as above.

### Quantification and Statistical Analysis

Statistical analyses were performed with Student t-test using Prism 10. All errors correspond to the standard deviation of the population. Statistically significant differences are indicated with four stars indicating an “extremely significant” result, corresponding to p ≤ 0.0001 while no stars typically means “not significant” (ns), which corresponds to p > 0.05. The number of samples analysed for each experiment is indicated in figure or table legends.

## Acknowledgements

We thank Daniel Abbühl for advice on tubulin experiments, Aline Alves for training on IFT and motility analyses and Amaia Ochandorena Saa and Sigolène Meilhac for stimulating discussions that led to the initiation of this project. We thank Daniel Abbühl, Serge Bonnefoy and Brice Rotureau for critical reading of the manuscript. We are grateful to the Photonic Bioimaging and to the Ultrastructural Bioimaging facilities for access to their equipment. We are thankful for the support of the UBI equipment from the French Government Programme Investissements d’Avenir France BioImaging (FBI, N° ANR-10-INSB-04-01).

This work was supported by La Fondation pour la Recherche Médicale (EQU202203014654), the ANR (ANR-18-CE13-0014-01) and a French Government Investissement d’Avenir programme, Laboratoire d’Excellence “Integrative Biology of Emerging Infectious Diseases” (ANR-10-LABX-62-IBEID). This work was supported by the Swiss State Secretariat for Education, Research and Innovation (SERI) contract MB22.00075 attributed to PG.

The authors declare that they have no conflict of interest.

## Author contributions

C. Girard-Blanc: Investigation, Formal analysis, Methodology, Validation, Visualisation, Writing - review & editing, T. Blisnick: Investigation, Validation, Visualisation, Writing - review & editing; V. Louvel: Investigation, Methodology; P. Guichard: Supervision, Writing - review & editing.; V. Hamel: Supervision, Writing - review & editing; P. Bastin: Conceptualization, Funding acquisition, Supervision, Writing - Original Draft, Writing - review & editing.

---

**Figure S1.**
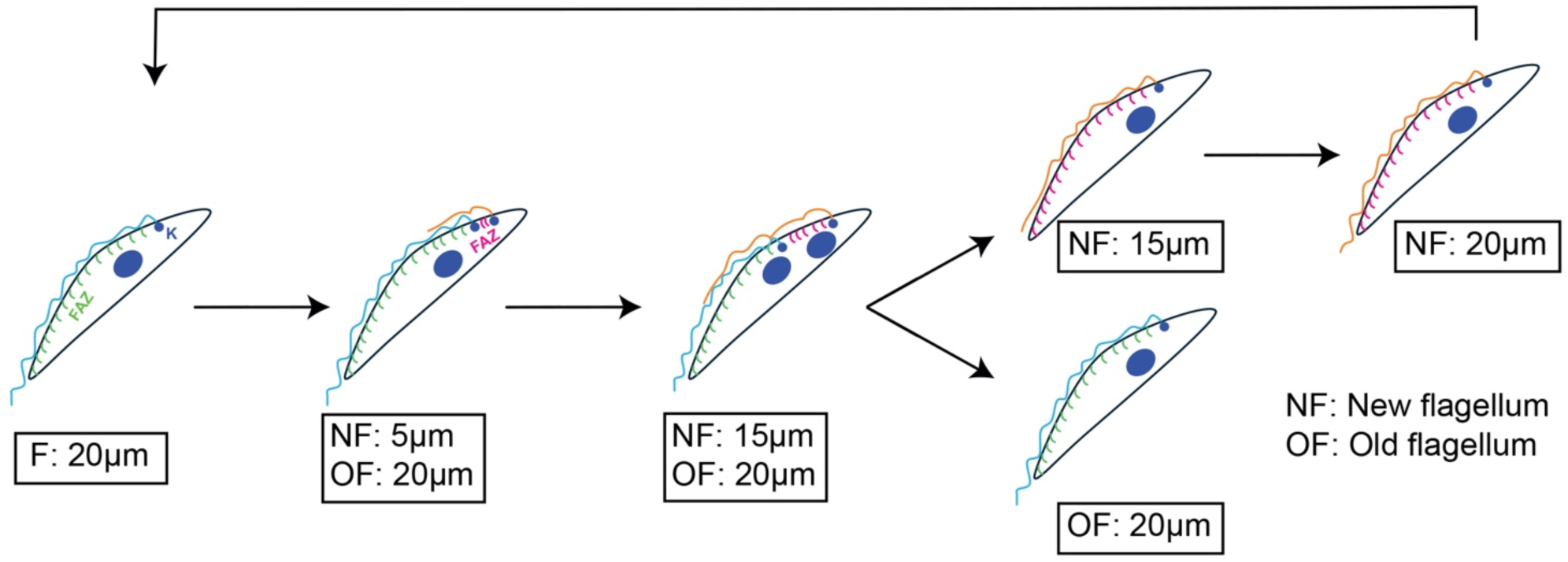
*T. brucei* cell cycle. A trypanosome in G1 possesses a single flagellum (cyan) of ∼20 µm, one nucleus and one kinetoplast (both shown in blue). The new flagellum (orange) elongates while the kinetoplast duplicates and segregates, leading to a cell with two kinetoplasts and one nucleus. After mitosis, the cell contains two nuclei and two kinetoplasts (2K2N) and the flagellum has grown to about 80% of its final length. One daughter cell inherits the old flagellum, and the other one inherits the new flagellum after division. This one continues growing until it reaches the final length of 20 µm. Reproduced from (Bertiaux et al., 2018b).

**Figure S2.**
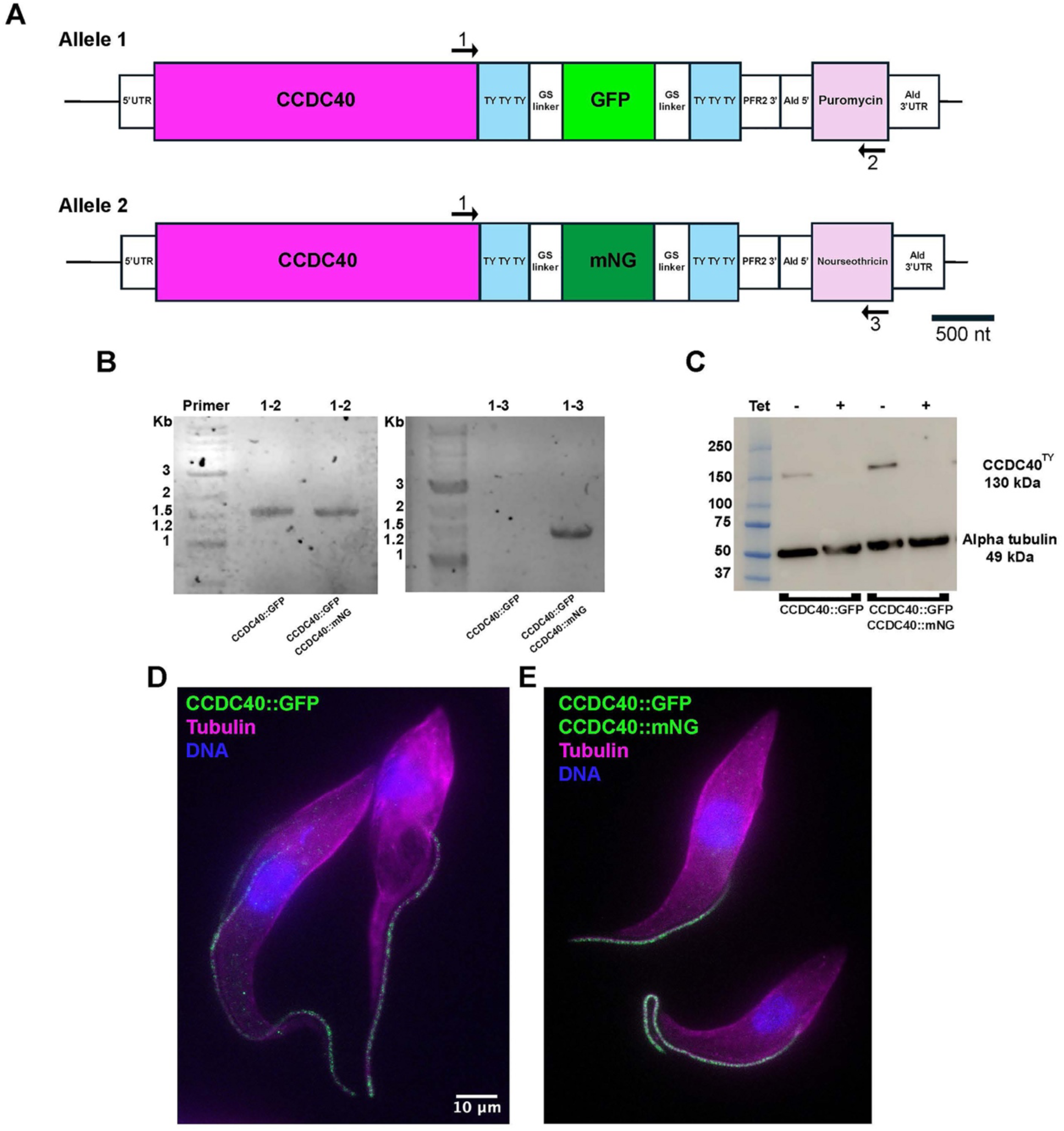
Tagging one or both CCDC40 alleles reveals flagellar positioning of the corresponding CCDC40::GFP protein. (A) Cartoon illustrating the integrated PCR product allowing tagging of CCDC40 with GFP or mNG and 6 Ty-1 epitopes, with repeats of glycine-serine as linkers (Dean et al., 2015). The short Ty-1 tags (10 amino acids each) and GS linkers (20 amino acids each) are not drawn to scale. The position of primers used to confirm proper integration is indicated and their sequence can be found in Table S1. (B) PCR analysis with the indicated pair of primers confirms proper integration of the construct upon GFP tagging or GFP and mNG tagging as indicated. (C) Western blot analysis of samples obtained from *CCDC40^RNAi1-500^* cell lines expressing either CCDC40::GFP, or both CCDC40::GFP and CCDC::mNG, grown without (-) or with (+) tetracycline for 48h, probed with BB2 to detect the tagged version and with TAT-1 for alpha-tubulin as loading control. The tagged protein is only detected in control conditions and effectively silenced upon RNAi. As expected, a stronger signal is observed in the double-tagged cell line. (D-E) 4-fold UExM of the same cell lines stained with alpha- and beta-tubulin and with BB2 to visualise the tagged version of CCDC40. On the side views (D), the signal looks more homogenous on the two-allele tagged cell line (E).

**Figure S3.**
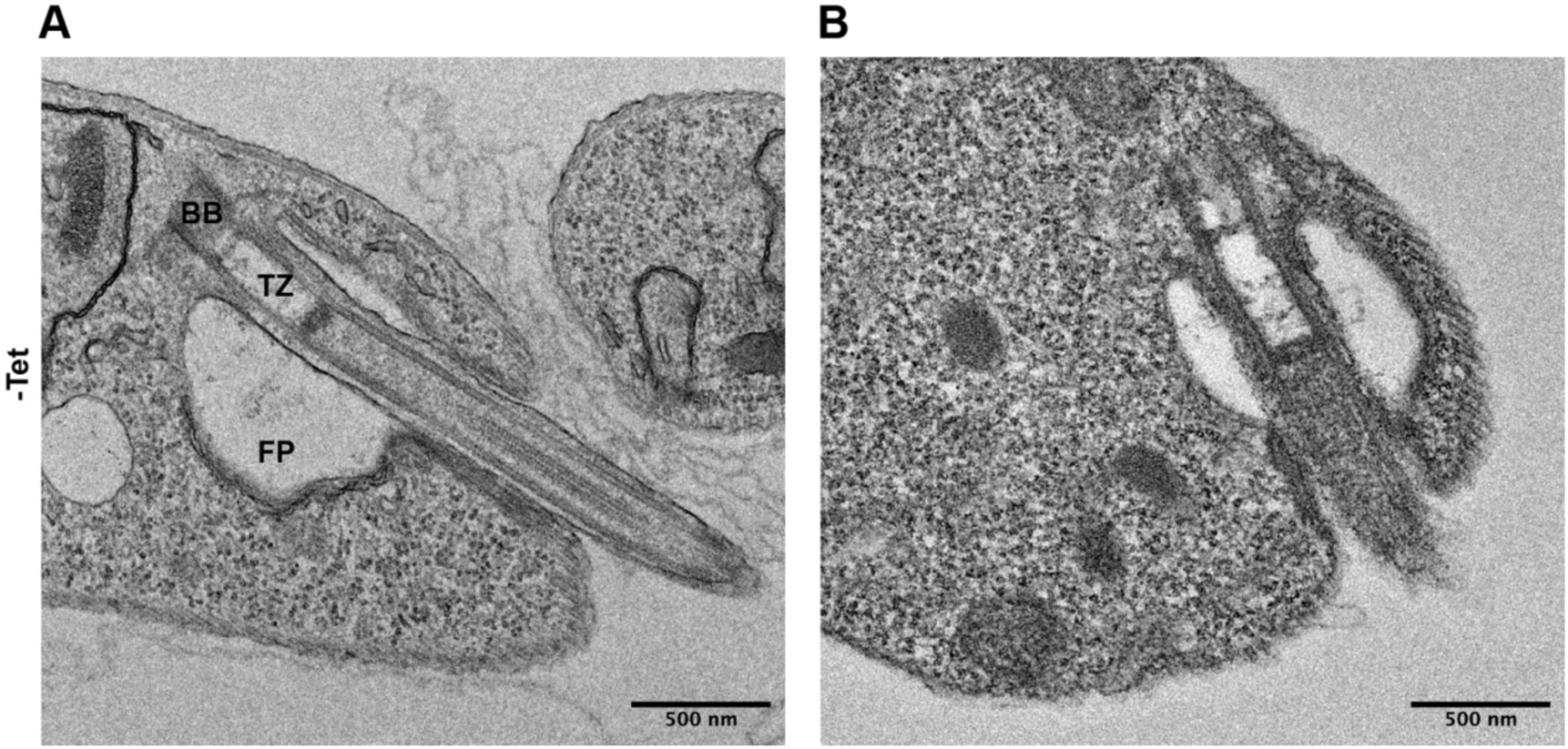
The proximal portion of the flagellum and the flagellar pocket remain intact upon depletion of CCDC40. *CCDC40^RNAi1100-1599^* cell line expressing CCDC40::GFP (single-allele tagged) were grown without (A) or with (B) tetracycline for 48h. Longitudinal sections sections of the base of the flagellum and flagellar pocket viewed by transmission electron microscopy, showing that the basal body (BB), the transition zone (TZ) and the flagellar pocket (FP) are normal in both conditions.

**Figure S4.**
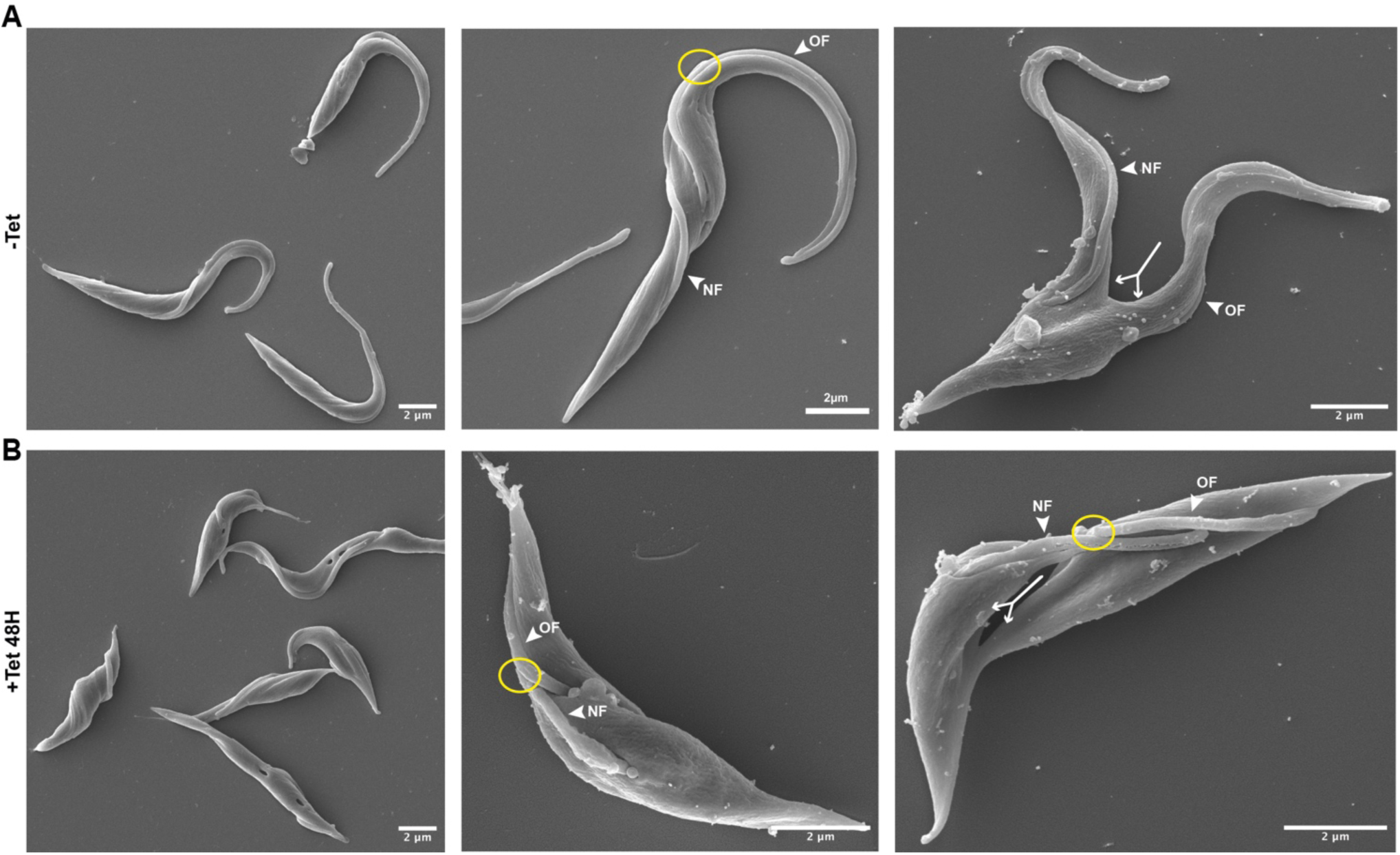
Knockdown of CCDC40 alters flagellum length and cell size. Scanning electron microscopy of the *CCDC40^RNAi1100-1599^* cell line expressing CCDC40::GFP (single-allele tagged) grown without (-Tet, A) or with (+Tet, B) tetracycline for 48h. Left panels show fields with several trypanosomes, the reduced length of both the cell body and the flagellum is visible. The central panels show cells growing a new flagellum (NF) that is connected to the old one (OF) by the flagella connector (yellow circle). Finally, the right panels show trypanosomes undergoing cell division with a clearly visible cytokinesis axis (arrowhead).

**Figure S5.**
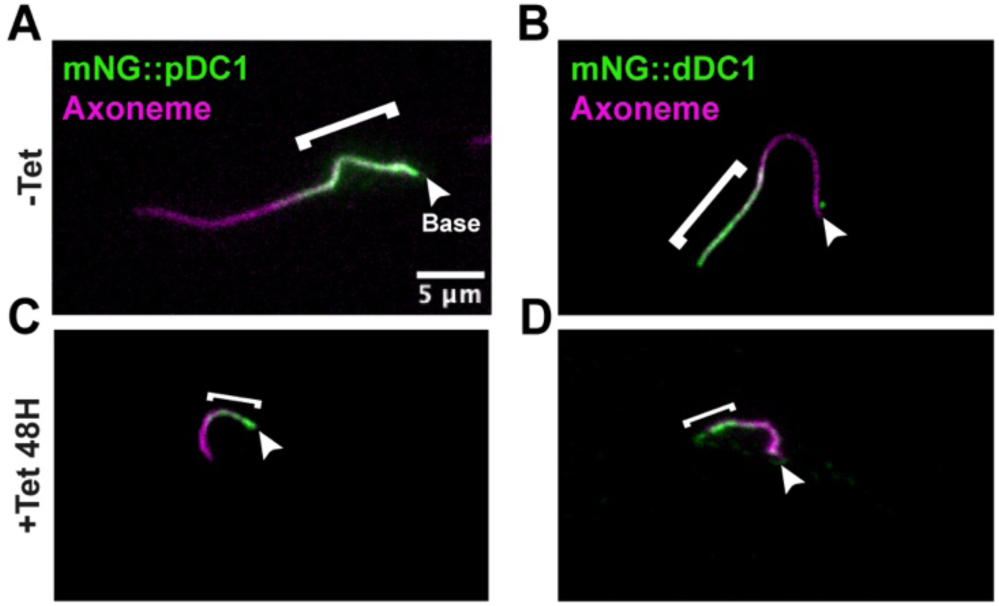
The distribution of the proximal and distal docking complexes is normal in the absence of CCDC40. *CCDC40^RNAi1-500^* cells expressing mNG::pDC1 (A) or mNG::dDC1 (B) were grown in absence (-Tet, top panels) or presence (+Tet, bottom) of tetracycline for 48h and stained with the axoneme marker mAb25 (magenta) and BB2 (green) for either tagged proximal (A,C) or distal (B,D) docking complex component. Independently of the stage of the cell cycle and of flagellar length, the proximal and distal docking complexes (indicated with white brackets) each spread for about half of the axoneme in both non-induced and induced conditions.

**Video S1. Knockdown of CCDC40 leads to profound alteration in microtubule organisation in the axoneme.** Full volume of *CCDC40^RNAi1100-1599^*cell line expressing CCDC40::GFP (single-allele tagged, green) cells grown without (-) or with (+) tetracycline for 48h, expanded 13-times by It-UExM and stained with the polyE antibody that detected tubulin.

**Video S2. Motility of cells depleted of CCDC40 is severely reduced**. *CCDC40^RNAi1-500^* cells were grown in absence (-Tet) or presence (+Tet) of tetracycline for 48h, imaged at low magnification (10-fold, left panels) for a total of 512 frames captured at 200 ms intervals. Videos were acquired with a 2 ms exposure time. High magnification is shown on the right panels.

**Video S3. IFT proteins still traffic on disrupted microtubules in flagella deprived of CCDC40.** *CCDC40^RNAi1-500^* cells expressing mNG::IFT81 as IFT reporter (Bertiaux et al., 2018a) were grown in absence (-Tet) or presence (+Tet) of tetracycline and videos were acquired for 30 seconds with a 100 msexposure time. Fields are shown on the left panels and zoom-in on an individual cell on the right. Robust IFT is visible in both control and induced cells, including in cases where microtubules are obviously disrupted.

## Notes

### Competing Interest Statement

The authors have declared no competing interest.

